# Concurrent stem- and lineage-affiliated chromatin programs precede hematopoietic lineage restriction

**DOI:** 10.1101/2020.04.29.069062

**Authors:** Fatemeh Safi, Parashar Dhapola, Sarah Warsi, Eva Erlandsson, Ewa Sitnicka, David Bryder, Charlotta Böiers, Ram Krishna Thakur, Göran Karlsson

## Abstract

The emerging notion of hematopoietic stem- and progenitor cells (HSPCs) as a low-primed cloud without sharply demarcated gene expression programs raises the question on how cellular fate options emerge, and at which stem-like stage lineage priming is initiated. Here we investigated single-cell chromatin accessibility of Lineage^−^, cKit^+^, Sca1^+^ (LSK) HSPCs spanning the early differentiation landscape. Application of a signal-processing algorithm to detect transition points corresponding to massive alterations in accessibility of 571 transcription factor-motifs revealed a population of LSK FMS-like tyrosine kinase 3(Flt3)^int^CD9^high^ cells that concurrently display stem-like and lineage-affiliated chromatin signatures pointing to a simultaneous gain of both Lympho-Myeloid and Megakaryocyte-Erythroid programs. Molecularly and functionally, these cells position between stem cells and committed progenitors, display multi-lineage capacity in vitro and in vivo, but lack self-renewal activity. This integrative molecular analysis resolves the heterogeneity of cells along hematopoietic differentiation and permits investigation of chromatin-mediated transition between multipotency and lineage restriction.

## INTRODUCTION

Helmed by the presence of rare, deeply quiescent, asymmetrically dividing and long-term self-renewing HSCs at the apex, hematopoiesis is often portrayed as the paradigm of tissue renewal (Orkin and Zon 2008). Even though the ultimate destiny of hematopoietic stem cells (HSCs) is to generate the entire functional diversity and numerical abundance of mature blood- and immune cells, accumulating evidence suggest that the initial steps of differentiation include subtle molecular changes as HSCs lose their self-renewal capacity and gradually become committed to different lineage fates (Laurenti and Gottgens 2018). These findings raise the question of how cellular potency and fate-decisions slowly manifest during the generation and restriction of the earliest progenitors from HSCs.

The classical model of hematopoiesis envisions a hierarchy wherein long-term (LT)-HSCs give rise to multipotent progenitors (MPPs) that manifest multilineage differentiation without long-term marrow reconstitution upon transplantation (Reya et al. 2001; Morrison et al. 1997; Yang et al. 2005). The MPPs are in turn succeeded by distinct cell fates that arise via discrete and increasingly lineage-restricted myeloid or lymphoid progenitors. An elaborate immunophenotyping strategy has divided MPPs into MPP1/short-term HSCs (ST-HSCs), MPP2, MPP3 and MPP4 based on a step-wise gain of CD34, CD48 as well as loss of CD150 cell-surface expression (Wilson et al. 2008; Cabezas-Wallscheid et al. 2017). Curiously, despite retaining multipotency and long-term self-renewal, HSCs themselves can be biased in terms of lineage output (Yu et al. 2017; Grover et al. 2016), and it is generally accepted that subsequent commitment of MPPs to a certain lineage is preceded by the expression of lineage-restricted gene signatures, known as lineage priming. While most of the HSC and MPP1 cells are not primed, MPP2s, MPP3s, and MPP4s are MegE, myeloid, and lymphoid-primed respectively; notably, each of the MPPs includes a different fraction of cells primed for specific lineages (Rodriguez-Fraticelli et al. 2018). However, whether multi-lineage primed cells are indeed present in MPP2 to 4, or whether these compartments are simply an assortment of unilineage-primed cells remains to be determined.

A reappraisal of cell potency, fate restriction, and heterogeneity has been invoked by single-cell analyses including transplantation, barcoding, lineage tracing and whole transcriptome sequencing (Cheng, Zheng, and Cheng 2020; Laurenti and Gottgens 2018). Single-cell RNA-sequencing (scRNA-seq) approaches, in particular, have advocated a continuum model of hematopoiesis wherein low-primed hematopoietic stem- and progenitor cells (HSPCs) give rise to unilineage-primed cells along the axes of differentiation (Velten et al. 2017; Giladi et al. 2018). These observations raise the question as to how cellular fate options emerge if the priming events at the level of gene expression are not sharply demarcated within HSPCs, and which cells, if any, begin to show frank lineage priming while retaining a partial stem-like program during the transition. It is tempting to speculate that lineage priming might instead initiate in the epigenome. By serving as an exquisite multi-layer template to mark gene regulatory regions in different states ranging from repressed, poised, and primed versus active, chromatin is an instructive template where lineage-priming is likely to be first manifested (Calo and Wysocka 2013; Bonifer and Cockerill 2017).

Here, we define the earliest stages of hematopoietic differentiation based on single-cell chromatin accessibility. Using high-throughput single-cell assay for transposase-accessible chromatin sequencing (scATACseq) approaches, we reveal distal regulatory regions and transcription factors uniquely active in HSCs and during lineage specification of either early megakaryocyte/erythroid (megE), or lymphoid/myeloid committed progenitors. Intriguingly, we identify an LSKFlt3^int^CD9^high^ population, where primitive and committed chromatin landscapes co-exist in individual cells representing a highly lineage-primed multipotent progenitor. Thus, our findings permit analysis of cellular and molecular events of a transition stage where self-renewal and multipotency is rapidly lost in favor of lineage commitment.

## RESULTS

### Single-cell chromatin analysis captures the heterogeneity and molecular transitions in hematopoietic stem cells and early progenitors

From LTHSC to MPPs, self-renewal potential is gradually lost with up-regulation of FMS-like tyrosine kinase 3 (Flt3) (Adolfsson et al. 2001; Christensen and Weissman 2001). It has been shown that Flt3^+^ multipotent progenitors serve as developmental intermediates for hematopoietic lineage specification (Boyer et al. 2011; Buza-Vidas et al. 2011; Akashi et al. 2000; Beaudin, Boyer, and Forsberg 2014; Kondo, Weissman, and Akashi 1997) where the 25% of LSK cells expressing the highest levels of Flt3 have been defined as restricted lymphoid-primed multipotent progenitors (LMPPs) (Adolfsson et al. 2005). As cells transition from LT-HSCs to LMPPs, MegE priming is down regulated and lymphoid priming is gained while granulocyte-monocyte potential is retained (Mansson et al. 2007). Thus, the earliest differentiation steps in hematopoiesis occur along the axis of Flt3 expression where multipotency is suggested to be largely restricted to the LSKFlt3^−^ compartment. This led us to postulate that progenitors with an intermediate level of Flt3 expression (Flt3^int^ henceforth) likely contain cells that manifest a transitional molecular program bridging multipotency with lineage commitment.

Recent observations provide compelling reasons to consider that lineage priming and fate changes in HSCs (Yu V et al. 2017) and early progenitors (Weinreb et al. 2020) is anchored in chromatin without stark manifestation at the gene expression level. Thus, to define cell states and underlying chromatin changes during the earliest stages of hematopoietic lineage priming, we analyzed 2,680 cells from immunophenotypically defined populations spanning early HSPCs (Table S1; Figure S1A-B) by droplet single-cell ATAC-sequencing (scATACseq) (see supplementary methods). The cells displayed a fragment size distribution that is characteristic of ATAC-assays, high enrichment at transcription start sites (TSS), an average of 25 × 10^3^ peak region cut sites fragments that mapped to the nuclear genome, with approximately 67% of Tn5 insertion sites detected within aggregate peaks from all samples (Fig. S2-S3). ScATAC-seq captures accessible chromatin regions in promoters as well as distal regulatory regions; therefore peak regions between −2000 to +500 bp relative to TSS were defined as ‘promoter/proximal’, while the peaks outside of promoters and gene bodies were denoted as ‘distal’. From a total of approximately 283,358 peaks obtained across all cells, we designated 107,011 as distal and 37,945 as promoter-proximal peaks while the rest mapped to gene bodies. Using Seurat’s graph clustering approach and uniform manifold approximation and projection (UMAP) (Becht et al. 2018), we identified and visualized 3 coarse, and 12 fine clusters of cells displaying distinctive global chromatin accessibility profiles (see supplementary methods; fine clusters described below), and annotated the clusters based on FACS-sorting markers to obtain classically recognizable stem and early progenitor populations such as LT and ST-HSC, MPP2, pre-MegE, and LMPP (Figure 1A-B).

**Figure 1.**
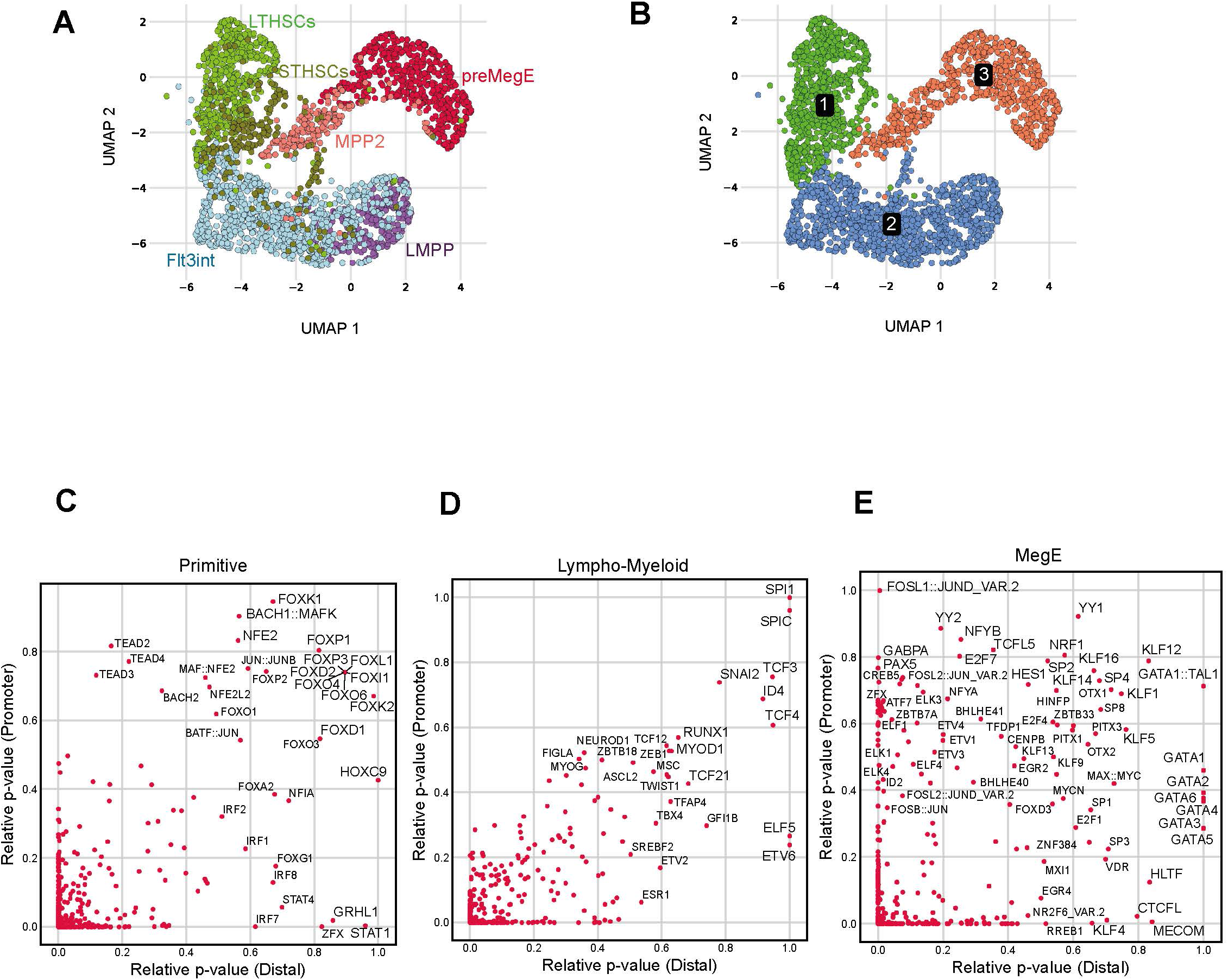
Chromatin accessibility captures the heterogeneity of early hematopoietic progenitors. UMAP embedding of cells with colours indicating **(A)** sorted HSPC populations, and **(B)** three coarse clusters displaying distinctive global chromatin accessibility profiles. Scatter plots indicating the relative p-value of TFBS enrichment in primitive **(C)**, Lympho-Myeloid **(D)** and MegE coarse clusters **(E)**. The relative p values for TFBS in distal- and promoter-proximal regions are depicted on the x-axis and y-axis respectively.

We next investigated whether there was any preferential accessibility of specific transcription factor (TF)-motifs as HSCs differentiate to either Lympho-Myeloid, or MegE lineages, and if the accessible motifs were localized in a specific genomic region (promoter-proximal or distal) (Figure 1 C-E, S4A-C). The primitive cells (relative to the other two coarse clusters) showed higher specific accessibility of the TEAD transcription factor family (TEAD1-4) motifs in promoter regions, while the motifs for established HSC regulators such as the FoxO family were enriched in both promoters and distal regions (Figure 1C). On the other hand, the motif for SPI1 (PU.1), a prominent myeloid and lymphoid lineage specifying TF (Fisher and Scott 1998; Bryder and Sigvardsson 2010) was enriched in both the promoters and distal peaks within cells of the Lympho-Myeloid lineage (Figure 1D). In comparison, GATA1-TAL1 (Wu et al. 2014) as well as KLF motifs (McConnell and Yang 2010) were most enriched at both promoters and distal peaks in the MegE cluster (Figure 1E). Together, the data suggest that the accessibility of regulatory chromatin regions for established key lineage-specifying TFs defines early hematopoietic cell type and lineage commitment.

### LSKFlt3^int^CD9^high^ cells concurrently display stem cell- and lineage-affiliated chromatin accessibility signatures

The two-dimensional UMAP layout of the scATACseq data shows that early commitment of HSPCs is associated with continuous rather than step-wise accessibility of the chromatin landscape, and suggests that the sparsely investigated LSKFlt3^int^ fraction represents a heterogeneous population containing cells with chromatin profiles spanning from multipotent progenitor-to LMPP-like (Figure 1A). Hence, we asked whether the LSKFlt3^int^ cells with more primitive chromatin landscapes could be prospectively isolated by virtue of retaining surface markers shared with stem cells. We have previously shown that cell surface expression of the tetraspanin CD9 marks all HSCs in murine bone marrow (Karlsson et al. 2013). CD9 has additionally been demonstrated to be incompatible with early Lympho-Myeloid differentiation and thus useful for purification of megakaryocyte progenitors (Nakorn, Miyamoto, and Weissman 2003). Moreover, increased CD9 protein-level coincided with enhanced megakaryocyte priming within HSCs (Nakamura-Ishizu et al. 2018). Thus, we investigated the relative abundance of CD9^+^ cells along the Flt3 expression spectrum by FACS analysis. Interestingly, we found that HSC-like cell-surface expression levels of CD9 could be detected in a small fraction of LSKFlt3^int^ cells, while LMPPs almost exclusively consist of CD9^low^ cells (Fig. 2A; S4D). Notably, only a minority of LSKFlt3^int^CD9^high^ cells co-expresses the stem cell marker CD150, which is in stark contrast to LSKFlt3^−^CD9^high^ HSCs (Figure S4D). Following scATACseq, prospectively isolated LSKFlt3^int^CD9^high^ and LSKFlt3^int^CD9^low^ cells were visualized onto the single-cell chromatin accessibility landscape of HSPCs. Intriguingly; the LSKFlt3^int^CD9^high^ cells (yellow) positioned close to the primitive LT- and ST-HSCs, while the LSKFlt3^int^CD9^low^ cells (dark blue) juxtaposed the LMPPs (Figure 2B; S4E). Of note, both populations occupied separate and non-overlapping positions within the LSKFlt3^int^ compartment signifying that CD9^high^ and CD9^low^ LSKFlt3^int^ cells represent two molecularly distinct populations.

**Figure 2.**
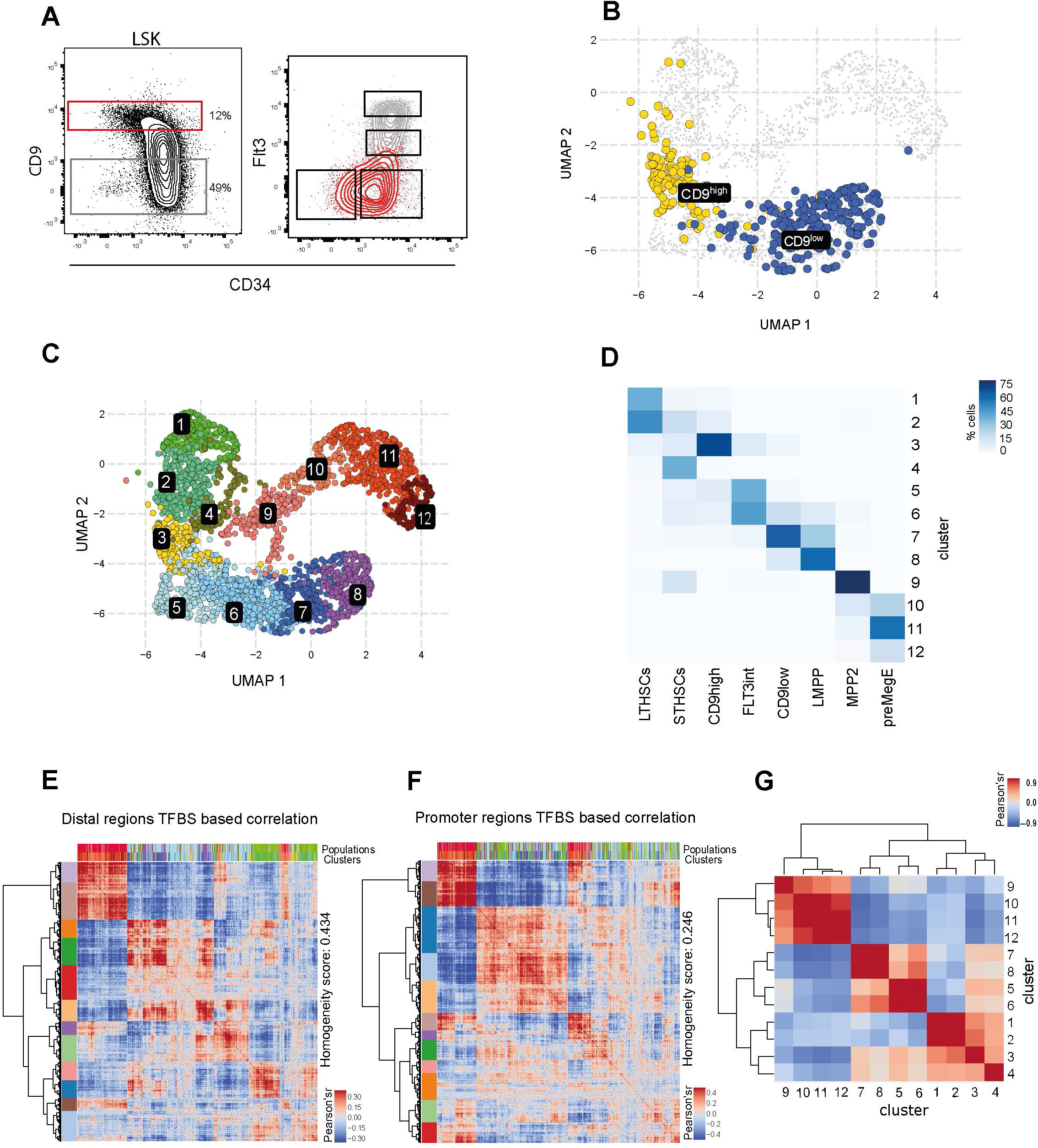
CD9 expression discriminates between LSKFlt3^int^ cells with distinct chromatin profiles. **(A)** Representative FACS plots showing the distribution of CD9^high^ cells within LSK HSPCs. **(B)** UMAP plot highlighting the position of LSKFlt3^int^CD9^high^ (yellow) and LSKFlt3^int^CD9^low^ cells (blue) based on chromatin accessibility relative to other HSPCs. **(C)** UMAP embedding of cells colour-coded based on 12 clusters identified in the data using Seurat. **(D)** Heat map showing the percentage of cells from each of the sorted population within a certain cluster. Darker colours indicate higher percentage. Heat maps showing cell-cell correlation based on 571 TFBS motifs detected in 107,011 distal peaks **(E)** or 37,945 promoter proximal peaks **(F)**. The color blocks along the rows indicate cell partitions obtained through hierarchical clustering of the cell-cell correlation matrix. **(G)** Cluster heat map showing correlation values (Pearson’s r-values) between clusters. Correlations were based on cumulative accessibility of TFBS motifs across distal regions.

Our dataset of single cells across prospectively isolated HSPCs permitted dissection of the chromatin basis for the observed continuum. Based on chromatin accessibility in both distal as well as promoter-proximal peaks, we identified and visualized 12 fine cell clusters (Figure 2C). Annotation of the FACS-sorted populations onto the chromatin clusters revealed that while each chromatin-defined cluster in general contained cells from one of the FACS-sorted groups, it additionally contained cells from other immunophenotype-sorted populations (Figure 2D). Interestingly, LT-HSCs contained two clusters (1 and 2) based on chromatin accessibility. While cluster 1 appears to be a distinct group with fewer cells (35% of LT-HSCs), cluster 2 is numerically abundant (>60% of LT-HSCs) and permeates over to the ST-HSCs. In comparison, the cells belonging to the classic ST-HSC/MPP1 immunophenotype displayed profound heterogeneity being present in six different clusters (2-7, and 9). Similarly, the LSKFlt3^int^ and Pre-MegE cell populations mapped to three or more chromatin clusters illustrating heterogeneity in terms of chromatin accessibility (Figure 2D).

To identify the importance of distal vs. proximal accessible regions for cell type specificity, we compared cell-cell correlations based on 571-transcription factor binding site (TFBS) motifs detected in either distal versus promoter-proximal peaks (Figure 2E-F, S4A-C). Interestingly, the distal peaks had higher discriminatory potential to separate cell clusters compared to the promoter-proximal peaks (homogeneity score: 0.434 vs. 0.246 respectively) indicating that the accessibility of distal regulatory regions is more likely to uniquely define a cell type. The correlation analysis based on distal peaks revealed that while LSKFlt3^int^CD9^high^ cells (cluster 3) have the highest chromatin similarity to ST-and LT-HSCs (clusters 1,2, and 4); the majority of CD9^low^ cells (cluster 7) correlated strongly with LMPPs in cluster 8 (Figure 2G). Collectively, this analysis suggests that global chromatin accessibility profiles transcend across immunophenotypically distinct populations and that CD9 could discriminate between cells with different chromatin landscape within the heterogeneous LSKFlt3^int^ population. Despite the continuum of chromatin accessibility observed in the scATACseq profiles, it is well established that lineage restriction occurs at some point between the LSKFlt3^−^ stem cells and the LSKFlt3^high^ LMPPs. However, at which stage the transition from multipotency to lineage restriction occurs, and how it is reflected on chromatin level remains unknown. To address this, we used the scATACseq data to identify transition points among the HSPC chromatin continuum by locating cells that display large changes in accessibility, and consequently major shifts in potency and lineage priming. To this end, we adopted a combinatorial strategy wherein cells were arranged along a lineage differentiation trajectory according to pseudotime, and TFBS motif accessibility were identified within the distal open chromatin regions of cells along the trajectory. Subsequently, the cells were analyzed for transitions in motif accessibility by aggregating changes across the compendium of 571 TFBS profiles (see supplementary methods; Fig. 3A-D and Fig. 3F-I).

**Figure 3.**
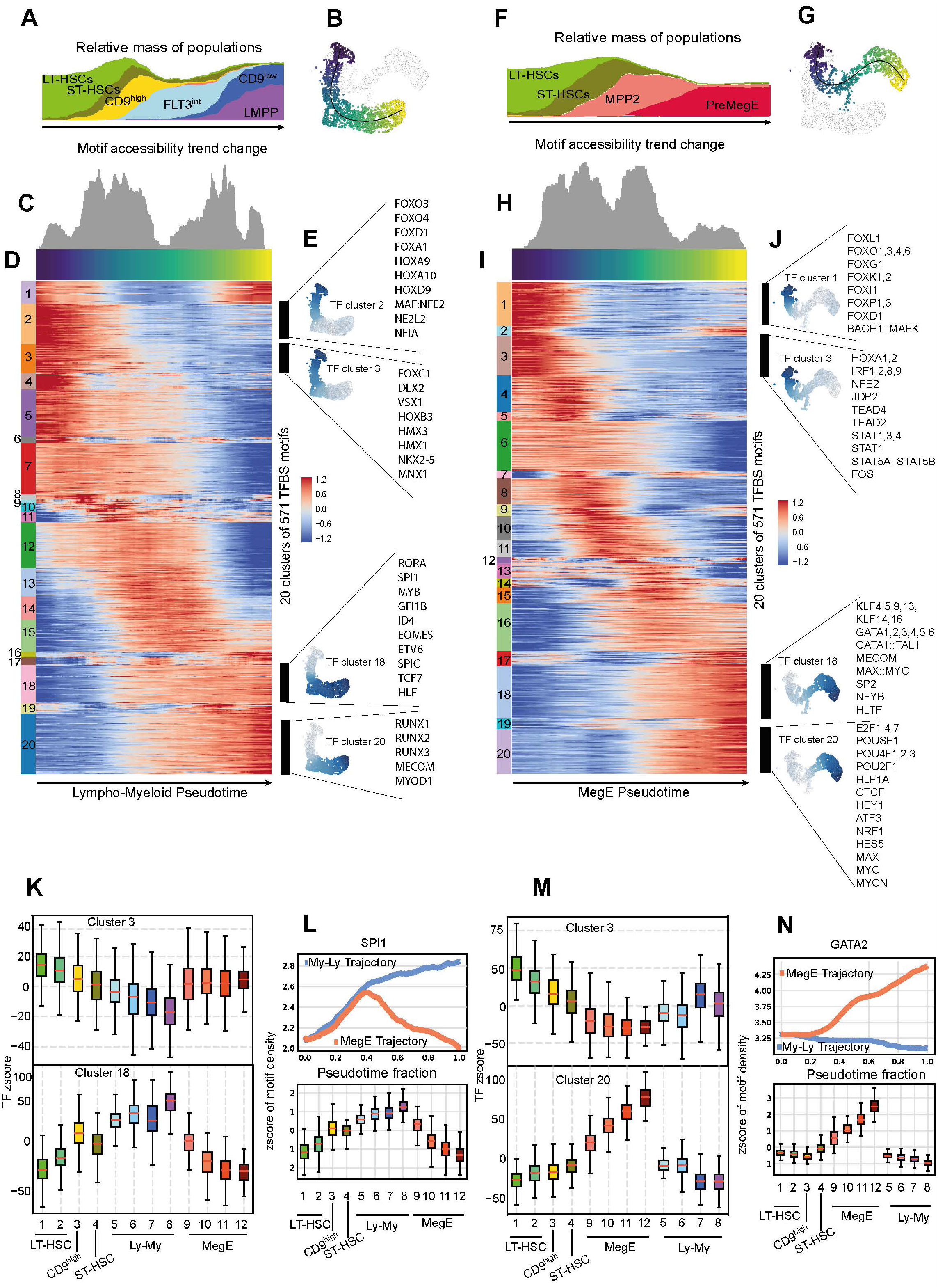
Dynamics of transcription factor motif accessibility during transition from stem cells to lineage specific progenitors. **(A**) Relative density of cells from sorted populations along the **(B)** Lympho-Myeloid trajectory as highlighted in the UMAP plot. The dark blue colour represents the early pseudotime while light yellow shows late pseudotime. **(C)** Area plot showing the frequency of trend change points along the Lympho-Myeloid trajectory. **(D)** Heat map displaying the relative accessibility of TFBSs across the Lympho-Myeloid trajectory. The cells are ordered according to the pseudotime on the x-axis and the rows represents single TFBS motif. The clusters of TFBS accessibility patterns are shown on the left of the heat map with TF-cluster numbers indicated. **(E)** Cumulative accessibility of representative TFBSs in TF-cluster 2, 3, 18 and 20 of the Lympho-Myeloid trajectory visualized on the UMAPs. **(F)** Relative density of cells from sorted populations along **(G)** the MegE trajectory as highlighted in the UMAP plot. **(H)** Area plot showing the frequency of trend change points along the MegE trajectory. **(I)** Heat-map showing the relative accessibility of TFBSs across the MegE trajectory. The cells are ordered according to the pseudotime on the x-axis and the rows represents single TFBS motif. The clusters of TFBS accessibility patterns are shown on the left of the heatmap with TF-cluster numbers indicated. **(J)** Cumulative accessibility of representative TFBSs in cluster 1, 3, 18 and 20 of the MegE trajectory visualized on the UMAP plot of the MegE trajectory cells. **(K)** Distribution of TFBS accessibility of TFBSs in cluster 3 and cluster 18 from Lympho-Myeloid trajectory. **(L)** SPI1 motif accessibility along Lympho-Myeloid and MegE trajectory. The panel below shows distribution of SPI1 accessibility in cells from each cluster. **(M)** Distribution of TFBS accessibility in cluster 3 and cluster 20 from the MegE trajectory. **(N)** GATA2 motif accessibility along Lympho-Myeloid and MegE trajectory. Distribution of GATA2 accessibility in cells from each of the clusters is shown in the bottom panel.

This approach resulted in identification of clusters of transcription factor (TF) binding site motifs that showed concerted changes reflecting either increased, decreased, or, oscillating accessibility along the pseudotime from LT-HSCs to different lineages. Notable examples included motifs of Hox and FoxO family from TF-clusters 2 and 3 that were highly accessible in primitive cells, but showed either strong reduction (HOXA9-10), or diminished accessibility (e.g. FoxO1) along the Lympho-Myeloid axis (Figure 3E; top panel; Table S3). In contrast, motifs of TF-clusters 18 and 20 exhibited higher accessibility in the Lympho-Myeloid committed progenitors compared to the primitive cells. These included previously known regulators like SPI1 (PU.1), RUNX1, MYB, and GFI1A/B (Figure 3E; bottom panel) (Bryder and Sigvardsson 2010, Ichikawa M, 2013, Wang X 2018). Similar analyses along the HSC-MPP2-preMegE pseudotime axis revealed increased accessibility of motifs for TFs previously implicated in MegE differentiation including KLF4, the GATA family, and TAL1 (Figure 3 F-J; Table S4) (CS Park 2016, Tremblay 2018, Vagapova 2018). A close inspection of trend changes along the HSC to Lympho-Myeloid pseudotime revealed areas of large changes in TF motif accessibility within certain cell populations (Figure 3C). Strikingly the majority of cells making up the earliest transition point mapped to cell-cluster 3, dominated by LSKFlt3^int^CD9^high^ cells while the other transition point occurred in LSKFlt3^int^CD9^low^ cells and LMPPs (Figure 3A).

In parallel, simultaneous examination of TF motif accessibility (TF-clusters 1-20) within cell-clusters 1-12 showed that motifs of TF-cluster 3 displayed progressively diminishing accessibility along the cell-clusters 1-8 as stem cells differentiate to Lympho-Myeloid progenitors (Figure 3K). In contrast, Lympho-Myeloid lineage TFs belonging to TF-cluster 18 showed a gradient of low to increasingly higher accessibility along cell-clusters 1-8. Intriguingly, the LSKFlt3^int^CD9^high^-dominated cell-cluster 3 exhibited unique concurrent accessibility of stem-like and lineage specific TF motifs within single cells (e.g FoxO, Hox-family and Spi1, Etv6, Runx-family, and Mecom respectively) indicating a state of global lineage priming within LSKFlt3^int^CD9^high^ cells (Figure 3D and 3K). Moreover, cell-clusters 9-12 affiliated with MPP2 and the pre-MegE lineage did not display a gain of Lympho-Myeloid lineage TFs such as from TF-cluster 18 implying that lineage segregation is firmly in place in these cells (Figure 3K; bottom panel). For example, Spi1 motif accessibility increased along the Lympho-Myeloid axis especially in cell-clusters 7-8, while it diminished in cells along the MPP2-preMegE pseudotime indicative of a bifurcation in the usage of TF (Figure 3L; top and bottom panels). Similarly, GATA2 motif accessibility increased along the MegE axis while it declined along the Lympho-Myeloid axis (Figure 3M and 3N).

For pseudotime analysis, cell-cluster 1 was assigned as a supervised trajectory origin and minimum spanning tree from this cluster was fitted to the two terminal clusters (cell-cluster 8 and cell-cluster 12, selected by Slingshot in an unsupervised manner). In this process, it is important to note that multi-potent cells downstream of HSCs may have been assigned to only one lineage in this analysis. Curiously, the LSKFlt3^int^CD9^high^ cells were apposed to LT-HSCs in terms of accessibility of MegE motifs from TF-cluster 20 (Figure 3M; bottom panel) such as GATA2 (Figure 3N) suggesting that these cells still carry MegE-potential.

Motivated by the recent reports demonstrating that enhancers are transcribed in a cell-type specific manner (Andersson et al. 2014), we identified 12,739 scATAC-seq distal peak regions that overlapped with transcribed FANTOM5 enhancers (See supplementary methods). Subsequently, we visualized the co-accessibility patterns of the enhancers along lympho-myeloid and MegE axes, and divided these into 20 clusters. Moreover, we queried each enhancer cluster for the presence of enriched TFBS motifs.

Analysis of enhancers along the lineage pseudotimes showed that Cluster 1, 3 and 4 along the lympho-myeloid axis (Fig 4A), and clusters 1, 2 and 3 in MegE pseudotime (Fig. 4D) contain enhancers that are specifically accessible within HSCs and early multipotent cells. Next we identified early lineage priming along the lympho-myeloid and MegE axes by considering enhancers that showed relative gain of accessibility immediately downstream of HSCs. Interestingly, in the lympho-myeloid trajectory (Fig. 4A-B), cluster 15 (n=1289) contained early-lineage accessible enhancers that dramatically increased accessibility in cells of cluster 3 that harbour Flt3^int^CD9^high^ cells. Intriguingly, the most enriched TFBS motifs in these enhancers are established lineage-affiliated master regulators like SPI1, SPIC, RUNX1, and ETV6 (Fig. 4C). Similar analysis along the MegE trajectory (Fig. 4D-E) revealed that cluster 16 (n=1024) comprises of early-lineage accessible enhancers. Again, gain of accessibility of these enhancers relative to LT-HSCs was initiated in the Flt3^int^CD9^high^-enriched cell cluster 3, and continued further onwards saturating at cell cluster 10 (Fig. 4E), suggesting a role for these enhancers in early MegE priming. Similar to the lympho-myeloid trajectory these enhancers are enriched for motifs of established lineage-specifying TFs like the GATA family TFBS as well as CTCF (Fig. 4F).

**Figure 4.**
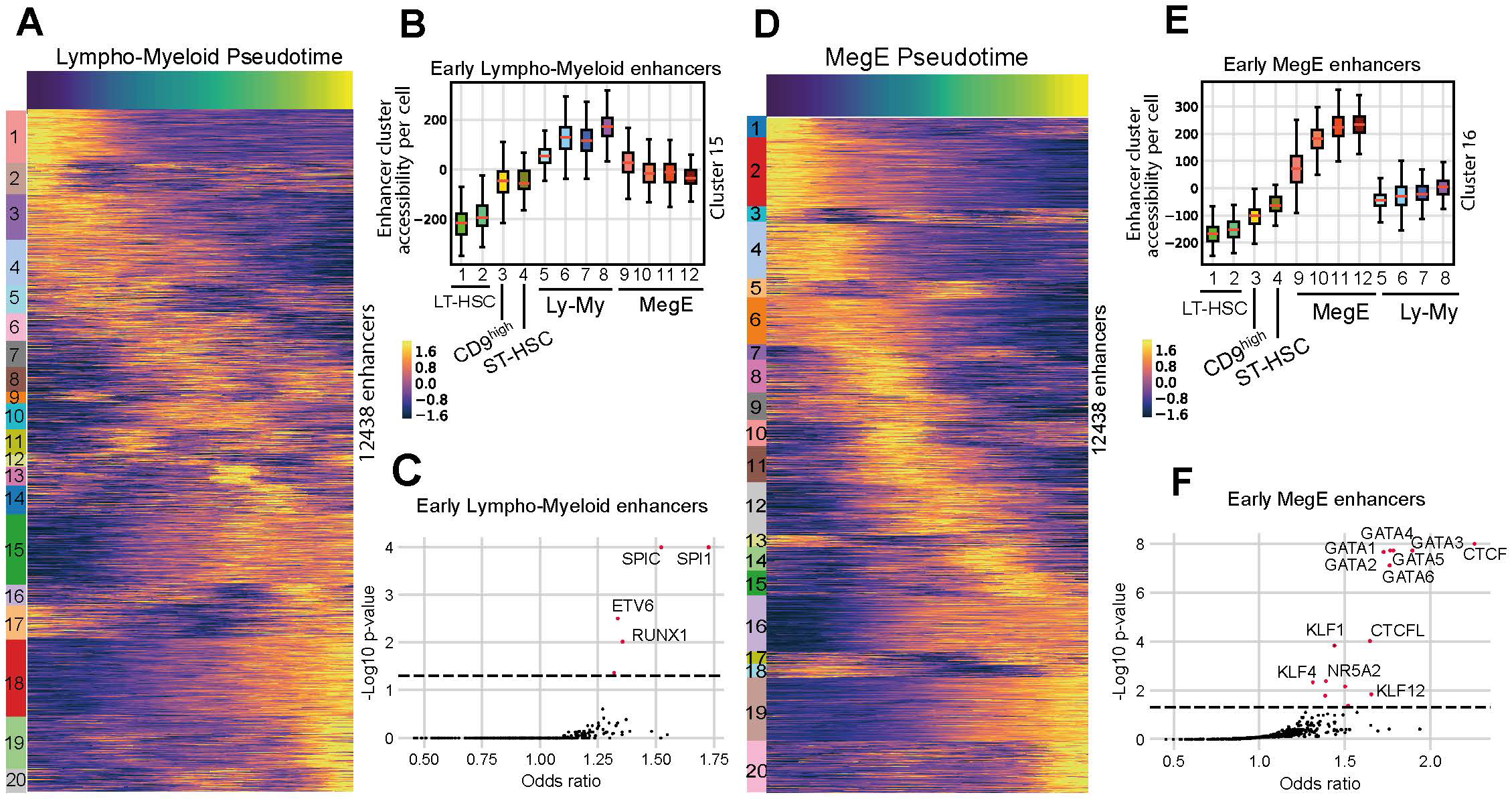
Dynamic accessibility of enhancers during early lineage specification. **(A)** Heat map showing relative accessibility scores (z scores) of individual enhancer peaks along the lympho-myeloid trajectory. The enhancer clusters obtained through hierarchical clustering are shown on the left bar with enhancer-cluster numbers indicated. **(B)** Boxplots showing the distribution of cumulative enhancer accessibility from enhancer-cluster 15 across each cell cluster. **(C)** Scatter plot showing enrichment of TFBS motifs in enhancer peaks from lymph-myeloid enhancer-cluster 15 compared to the rest of enhancers. **(D)** Heat map showing relative accessibility scores (z scores) of individual enhancer peaks along MegE trajectory. The clusters obtained through hierarchical clustering are shown on the left bar with cluster numbers indicated. **(E)** Boxplots showing the distribution of cumulative enhancer accessibility from enhancer-cluster 16 across each cell cluster. **(F)** Scatter plot showing enrichment of TFBS motifs in enhancer peaks from MegE-enhancer cluster 16 compared to the rest of enhancers.

Collectively, these findings indicated that massive changes in TF motif accessibility take place during the transition from stem cells to lineage specific progenitors, with substantial changes reflected in the LSKFlt3^int^CD9^high^ cells. Importantly, individual cells at this transitional state simultaneously display a chromatin accessibility profile similar to HSCs, as well as gain of lineage-affiliated molecular programs associated with Lympho-Myeloid and MegE differentiation trajectories. Moreover, the enhancer-centric investigation suggests that relative to HSCs LSKFlt3^int^CD9^high^ cells gain simultaneous accessibility of both lympho-myeloid and megE enhancers, that display motifs of lineage-specific TFs. Together with distal scATAC-seq peak based analysis, these results affirm that while LSKFlt3^int^CD9^high^ cells retain HSC-like enhancer accessibility, these represent a transition stage where global early multi-lineage priming is initiated.

### Single-cell RNA-Sequencing validates a gradient of primitiveness in Flt3^+^ cells marked by the level of CD9 expression

To validate the findings from scATACseq we performed a targeted heterogeneity analysis on LSKFlt3^int^ cells. Single-cell RNA sequencing (scRNAseq) was performed on LSKFlt3^int^ cells, LSKFLT3^int^CD9^high^ cells, and LSKFLT3^int^CD9^low^ cells. In total, data was obtained from 1720, 523 and 219 single cells from the LSKFlt3^int^, LSKCD9^high^ and LSKCD9^low^ populations respectively, analyzed by Seurat, and visualized as a two dimensional UMAP (Tim Stuart1 et al. 2018), this identified ten clusters of cells (Figure 5A). While the LSKFlt3^int^ cells were distributed across the UMAP landscape, LSKFlt3^int^CD9^high^ and LSKFlt3^int^CD9^low^ cells showed preferential enrichment in certain clusters indicating that CD9 expression correlates with differences in mRNA composition (Figure 5B). To quantify the distribution of LSKFlt3^int^CD9^high^ and LSKFlt3^int^CD9^low^ cells to particular clusters, a cell projection analysis (see supplementary methods) was performed wherein LSKFlt3^int^CD9^high^ and LSKFlt3^int^CD9^low^ cells were individually projected over LSKFlt3^int^ cells (Figure 5C, 5E). Upon projection, each LSKFlt3^int^ cell obtained a score that indicates its similarity to the cells from the projected population. In the case of LSKFlt3^int^CD9^high^ cells, the median score was highest in Flt3^int^ cells of cluster 3 (Figure 5D) while for LSKFlt3^int^CD9^low^ cells, the highest median score was observed in cluster 1 (Figure 5F). Similarly, cluster 3 had the largest relative assignment of LSKFlt3^int^CD9^high^ cells (31.74%) while 40.18% of LSKFlt3^int^CD9^low^ cells projected to cluster 1 (Figure 5G). Moreover, the relative enrichment of LSKFlt3^int^CD9^high^ and LSKFlt3^int^CD9^low^ cells over Flt3^int^ cells was again highest in cluster 3 (2.59) and cluster 1 (1.79), respectively (Figure 5H).

**Figure 5.**
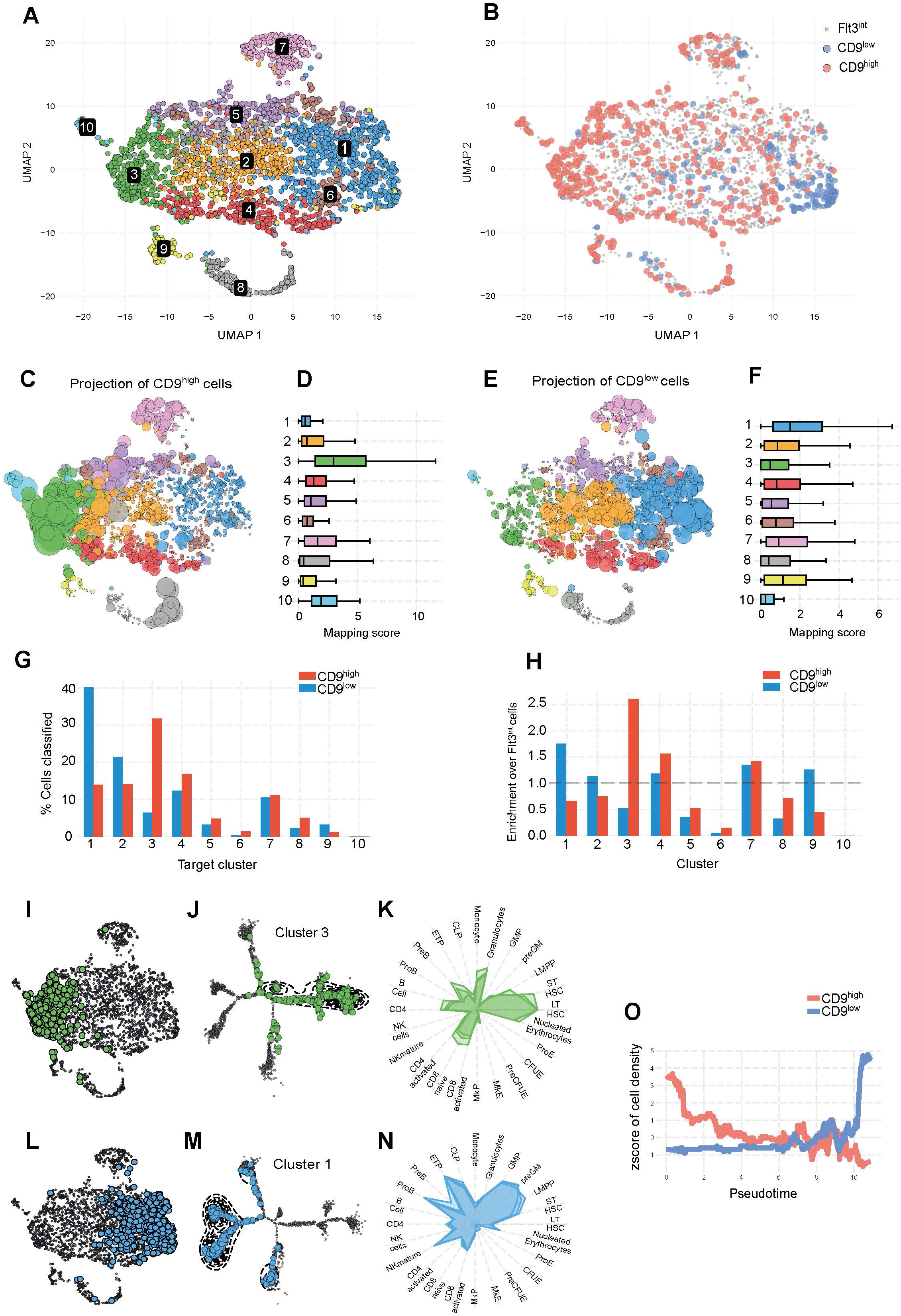
Single cell RNA-sequencing reveals a correlation between primitive mRNA signature and CD9 expression within the LSKFLT3^int^ compartment. **(A)** UMAP visualization of ten identified clusters (total 2462 cells) from pooled 1720 LSKFlt3^int^, 523 LSKFlt3^int^CD9^high^ and 219 LSKFlt3^int^CD9^low^ cells (**B).** Mapping scores obtained on projection of LSKFlt3^int^ CD9^high^ cells; visualized in UMAP of LSKFlt3^int^ cells **(C)**. The size of LSKFlt3^int^ cells indicates the magnitude of LSKFlt3^int^CD9^high^ sc mapping. Colour coding represents cluster identity. **(D)** Boxplots showing cluster-wise distribution of mapping scores obtained on projection of LSKFlt3^int^CD9^high^ cells. **(E)** Mapping scores obtained on projection of LSKFlt3^int^CD9^low^ cells visualized in UMAP layout of LSKFLT3^int^ cells. **(F)** Boxplots showing cluster-wise distribution of mapping scores obtained on projection of LSKFlt3^int^CD9^low^ cells. **(G)** Percentage of LSKFlt3^int^CD9^high^ and LSKFlt3^int^CD9^low^ cells assigned into each cluster. **(H)** Post-assignment enrichment of LSKFlt3^int^CD9^high^ and LSKFlt3^int^CD9^low^ cells in each cluster. Cells from cluster 3 highlighted in UMAP **(I)** and DDRTree **(J)** embedding of cells. **(K)** Radar plot showing cell type similarity of cluster 3 cells. Cells from cluster 1 highlighted in UMAP **(L)** and DDRTree **(M)** embedding of cells. **(N)** Radar plot showing cell type similarity of cluster 1 cells. **(O)** Density of cell populations along pseudotime. LSKFlt3^int^CD9^high^ cells population are highlighted in pink while LSKFlt3^int^CD9^low^ cells are highlighted in blue.

As the Flt3^int^ population contains a melange of actively differentiating progenitors, the cells were further arranged in a trajectory-using Monocle2 (Qiu et al. 2017). We found that cells from LSKFlt3^int^CD9^high^ and LSKFlt3^int^CD9^low^ populations clustered on divergent ends of the trajectory where cells from cluster 3 (Figure 5I) were exclusively present in the LSKFlt3^int^CD9^high^ enriched regions (Figure 5J). On the contrary, LSKFlt3^int^CD9^low^-enriched regions of the trajectory (Figure 4M) overlapped with cells from cluster 1 (Figure 5L). This confirmed our earlier observation that LSKFlt3^int^CD9^high^ and LSKFlt3^int^CD9^low^ populations do indeed represent distinct cell states within the LSKFlt3^int^ population. Furthermore, differentially expressed genes for each cell cluster were identified using Seurat (supplementary Figure S5A) and compared to available gene expression databases for various hematopoietic populations (Bagger et al. 2016). The resulting cell type specification for each cluster was visualized as radar plots. Cluster 3 showed enrichment in HSC-like signature (Figure 5K) as compared to other clusters (supplementary Figure S5B) indicating a primitive mRNA composition within LSKFlt3^int^CD9^high^ cells. In contrast, the molecular signature of the LSKFlt3^int^CD9^low^-enriched cluster 1 resembled that of LMPPs and pre-GMPs (Figure 5N). This was further supported by analysis of the density of cell populations along a pseudotime axis where cells from the LSKFlt3^int^CD9^high^ population accumulated early in comparison to the LSKFlt3^int^CD9^low^ cells (Figure 5O).

To directly assess molecular profile as a proxy for cell function we combined single-cell quantitative multiplex reverse transcriptase (RT)-PCR with index sorting for cell surface markers on 192 cells from the LSKCD34^+^Flt3^int^ population. For each index sorted single-cell, the cell surface expression of additional markers for primitive hematopoietic populations (CD48, CD150 (Yilmaz, Kiel, and Morrison 2006; Osawa et al. 1996) and CD9 (Karlsson et al. 2013) were recorded. The cells were analyzed by qRT-PCR against a panel of genes that included functional regulators and markers for HSC activity, lineage commitment, and cell cycle (Figure 6A and 6B; Table S2). Following unsupervised principal component analysis (PCA); five subpopulations with distinct molecular signatures could be identified highlighting the heterogeneity within the LSKFlt3^int^ population. By correlating each group to the gene signature, we observed one subpopulation with an HSC-like, quiescent signature (labeled in green) while the other subgroups are more proliferative. Moreover, the subpopulations marked by purple and blue express a MegE gene signature, while the red and yellow subgroups are more correlated to LMPP and the lymphoid signature (Figure 6A). Interestingly, we observed that *CD9* mRNA levels discriminate between subpopulations with HSC/MegE and LMPP signatures (Figure 6A). We next correlated these signatures with index sort data and found that CD9 expression (Fig. 6C) can capture the more primitive molecular signature (green) while neither subpopulations with LMPP-like signature nor those with a proliferative MegE signature expressed high levels of CD9 on the cell surface, despite the latter being *CD9^+^* at the mRNA level. As observed in previous FACS analysis, none of the subpopulations displayed cell-surface expression of the stem cell-marker CD150, while the low levels of cell-surface expression for the differentiation marker CD48 confirmed the primitive molecular signature observed in the CD9^+^ green subpopulation.

**Figure 6.**
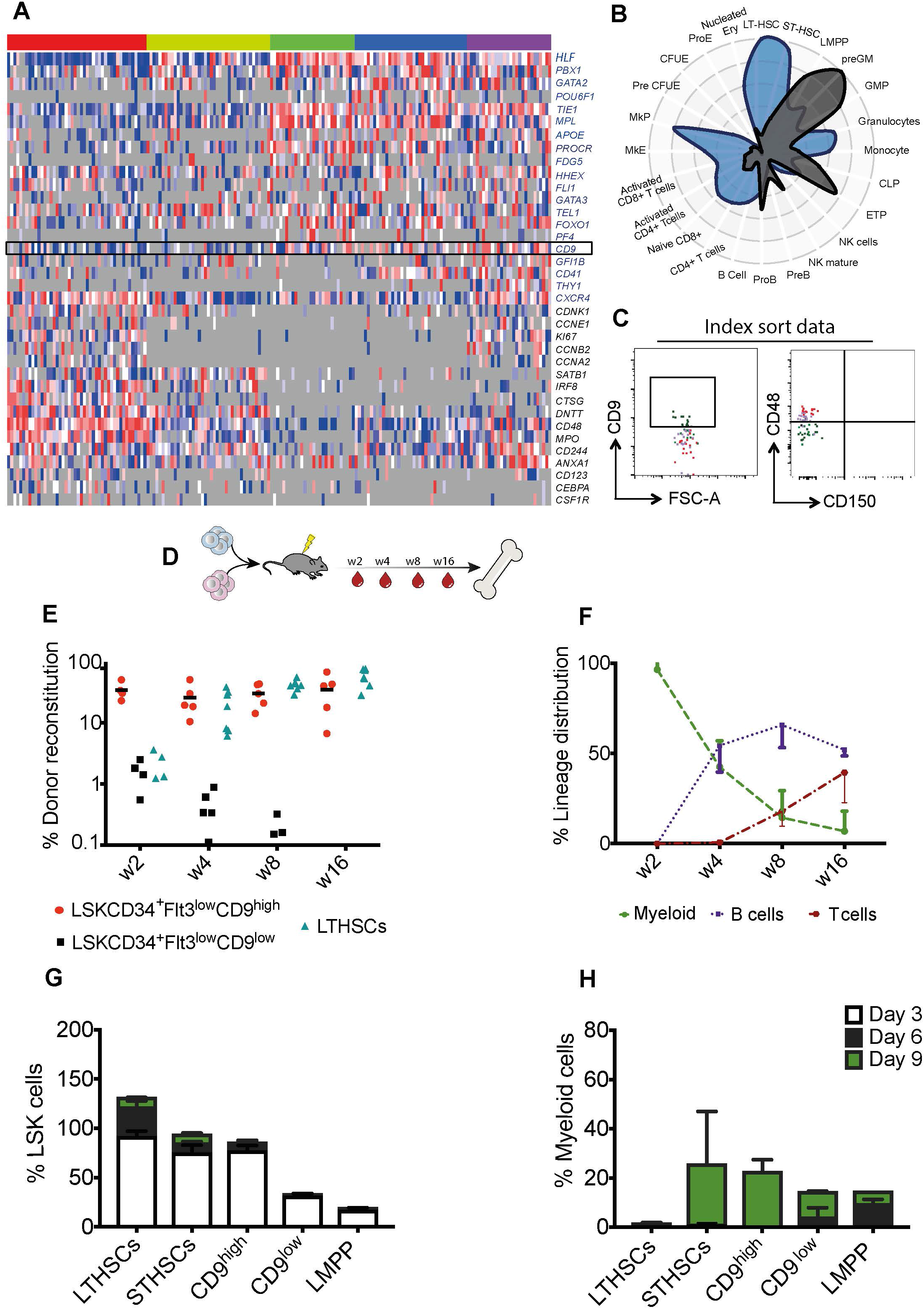
The LSKFlt3^int^CD9^high^ cells display multilineage potential but lacks long-term self-renewal capacity. **(A)** Fluidigm single-cell gene expression analysis on LSKFlt3^int^ cells presented as a heat-map with five molecularly distinct color-coded subpopulations. The level of gene expression is color-scaled with red indicating high- and blue low expression; gray denotes lack of- or undetected expression. Each row represents a gene primer and each column represents a single cell. **(B)** Radar plot of the genes with the most variable expression specifying the HSC/MegE clusters (blue) or Lympho-Myeloid clusters (grey) measuring enrichment of murine hematopoietic cell-type signatures. **(C)** Immunophenotypic characterization of the green, red and purple clusters from figure 2A using index sorting data. Each cell is color-coded according to the PCA-clustering of the single-cell gene expression analysis. **(D)** Experimental design for competitive transplantation experiments. **(E)** Donor reconstitution in peripheral blood in animals transplanted with 100 cells of the indicated populations. Each dot represents one recipient mouse (n=3), **(F)** Mean donor lineage distribution over time in recipients transplanted with 100 LSKCD34^+^Flt3^int^CD9^high^ cells. The error bars represent standard deviation (n=3). **(G-H)** Pooled data from FACS analysis of 500 cultured cells from the indicated populations, **(G)** Shows the fraction of LSK cells (n=3) and **(H)** the fraction of Gr1^+^/Mac1^+^ double positive cells at 3, 6 and 9 days of myeloid differentiation culture (n=2).

Thus, in accordance with the scATACseq data, scRNAseq analyses reveal that a fraction of Flt3^int^ expressing LSK cells are molecularly similar to primitive, multipotent progenitors, and that this molecular program correlates with the expression of CD9. Together, these results suggest that a transition point between hematopoietic multipotency and lineage restriction can be captured where cell-surface expression of CD9 and Flt3 is combined.

### LSKFlt3^int^CD9^high^ cells represent a multilineage hematopoietic transition state

To functionally validate the data from the sc-genomics experiments, we transplanted 100 LSKFlt3^int^CD9^high^ or LSKFlt3^int^CD9^low^ cells into irradiated recipient mice (Figure 6D). Interestingly, in contrast to the transient reconstitution observed from the LSKFlt3^int^CD9^low^ cells, long-term engraftment was observed in mice transplanted with LSKFlt3^int^CD9^high^ cells (Figure 6E and 6F). When transplanting limited doses (10 cells), LSKFlt3^int^CD34^+^CD9^high^ cells still reconstituted 5 out of 11 recipients (Supplementary Figure S6A). However long-term reconstitution was exclusive to the lymphoid lineages, while the myeloid cells were only observed during the first weeks following transplantation. Moreover, no long-term repopulation of the LSK compartment was detected, indicating that despite the observed multilineage engraftment, these cells lack self-renewal capacity (Figure S6B-S6C). Next, to establish the hierarchical relationship between CD9^high^ and CD9^low^ cells within the LSKFlt3^int^ population, we first performed short-term differentiation experiments in *vitro*, and tracked the changes in early stem and progenitor cell-surface markers: c-kit and Sca-1 by flow cytometry (Figure 6G and 6H) (Pietras et al. 2015). As expected, LT-HSCs and ST-HSCs showed persistent expression of LSK markers, whereas LMPPs rapidly lost the LSK profile and differentiated into c-Kit^+^Sca1^−^ and c-Kit^−^Sca-1^−^ myeloid progenitors (Supplementary Figure S7A-S7B). Within the LSKFlt3^low^ fraction, loss of the LSK cell-surface immunophenotype was substantially slower in CD9^high^ cells compared to the CD9^low^ cells, with a distinct LSK population still observed nine days after culturing the LSKFlt3^int^CD9^high^ cells, and a loss of LSK cells already at day six when starting with LSKFlt3^int^CD9^low^ cells (Figure 6G-H).

To address the clonal capacity of LSKFlt3^int^CD9^high^ cells *in vitro*, we designed a protocol to allow for single-cell differentiation into B, myeloid, or erythroid cells. Single-cells were plated directly onto OP9 stroma cells and following four days of co-culture, the proliferating clones were divided. Half of the progeny of each clone was transferred to erythroid culture conditions, and the other half was transferred to the B cell culture conditions (Figure 7A). Subsequently, the clones were analyzed by flow-cytometry for markers of erythroid (CD45^−^F4-80^−^Mac1^−^Gr1^−^TER119^+^), B (CD45^+^NK1.1^−^ CD19^+^), or myeloid cells (either CD45^+^ NK1.1^−^ Mac1^+^ Gr1^+^ or CD45^+^ NK1.1^−^ Mac1^+^ F-480^+^ (Figure 7B)). Strikingly, in these growth conditions allowing for several lineage decisions, 57% of the clone’s derived form LSKFlt3^int^CD9^high^ single-cells generated multilineage progeny (44% Myeloid-B and 13% Myeloid-Erythroid). In comparison, only 13% of clones of LSKFlt3^int^CD9^low^-origin consisted of several lineages (11% Myeloid-B and 2% Myeloid-Erythroid) (Figure 7C, 7D)). Together, this data provides functional support for the identification of an LSKFlt3^low^CD9^high^ multilineage-primed progenitor at the transition stage between stem and lineage committed cells.

**Figure 7.**
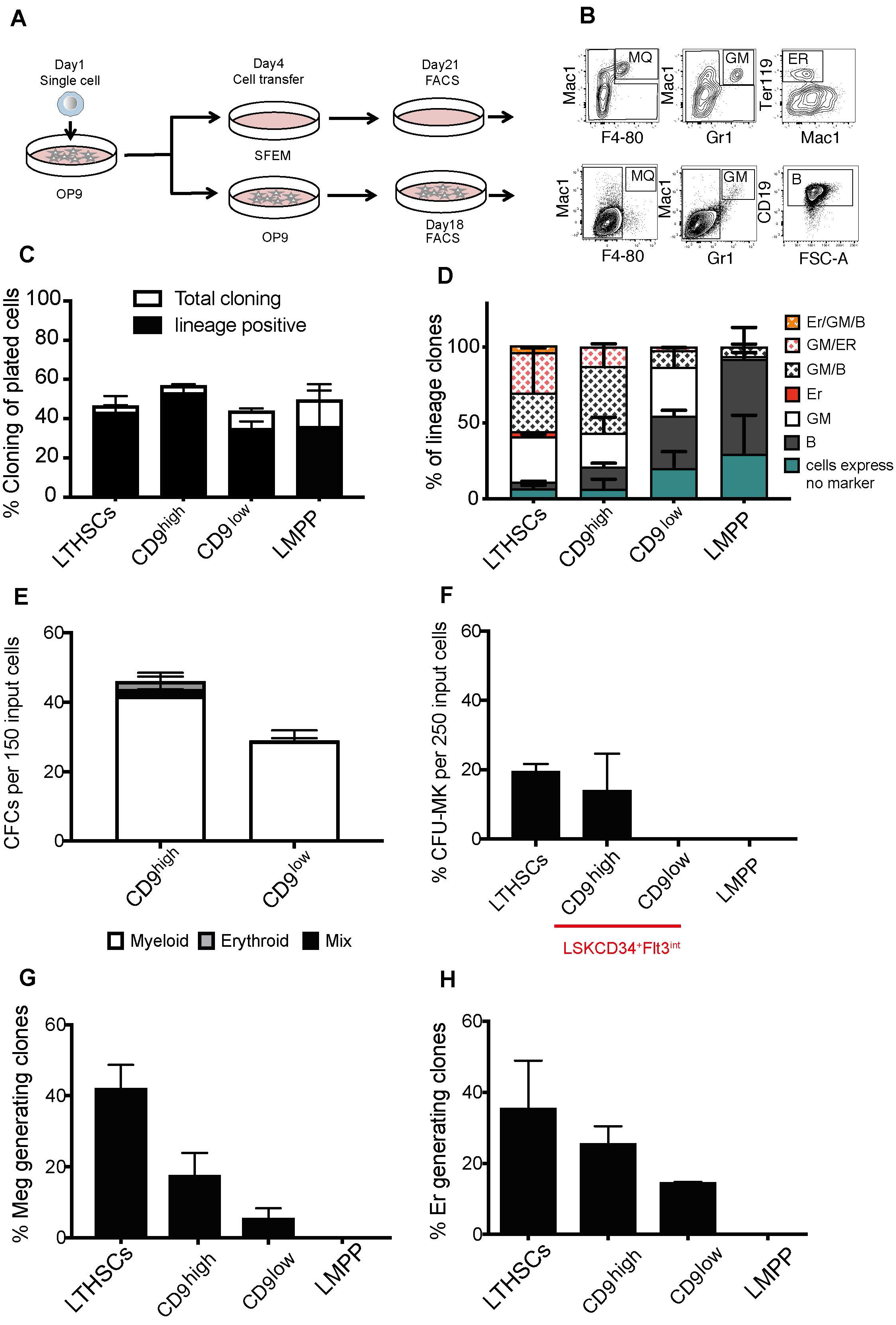
CD9 expression within LSKFlt3^+^ cells correlates with Meg/Erythroid differentiation potential. **(A)** Experimental strategy for clonal switch-culture experiments (n= 2) **(B)** Representative FACS profile of LSKFlt3^int^CD9^high^-derived GM/Macrophage/B/Erythroid clones. **(C)** Cloning frequency (white bar) and frequency of lineage positive clones (GM/Macrophage/B/E) (black bar). CD9^high^ = LSKFlt3^int^CD9^high^, CD9^low^= LSKFlt3^int^CD9^low^. **(D)** Lineage distribution of clones derived from single cells from the indicated sub-populations. CD9^high^ = LSKFlt3^int^CD9^high^, CD9^low^= LSKFlt3^int^CD9^low^ **(E)** CFC-capacity per 150 cells for different sub-populations following eight days of culture. (n = 3). Error bars indicate standard deviation. CD9^high^ = LSKFlt3^int^ CD9^high^, CD9^low^= LSKFlt3^int^CD9^low^ **(F)** Pooled data for CFU-Mk assay (n=2) CD9^high^ = LSKFlt3^int^CD9^high^, CD9^low^= LSKFlt3^int^CD9^low^. **(G)** Pooled data showing identified Megakaryocytes and **(H)** Erythroid cells following 10 days suspension culture of single cells from the indicated populations (n=3). Error bars represent standard deviation. CD9^high^ = LSKFlt3^int^CD9^high^, CD9^low^= LSKFlt3^int^CD9^low^.

The motif analysis of scATACseq data revealed an increase in MegE priming in LSKFlt3^int^CD9^high^ cells compared to LT-HSCs. Additionally, 13% of single LSKFlt3^int^CD9^high^ cells generated erythroid progeny in multilineage-supporting growth conditions. To further explore the MegE potential of these cells, we performed colony-forming cell (CFC) assays on the CD9^high^ and CD9^low^ LSKFlt3^+^ populations. In agreement with our previous data, CFC-forming capacity of LSKFlt3^int^CD9^high^ population was higher compared to the LSKFlt3^int^CD9^low^ cells. Additionally, even though the generation of BFU-E colonies was low, all erythroid potential associated with the LSKFlt3^int^CD9^high^ cells (Figure 7E).

To specifically measure megakaryocyte differentiation capacity, 500 cells from different LSK populations were seeded in colony-forming unit-megakaryocyte (CFU-MK) cultures (Berthier et al. 1993; Bruno, Briddell, and Hoffman 1988). Interestingly, megakaryocyte potential was exclusively observed in LT-HSCs and the LSKFlt3^int^CD9^high^ population, with neither LSKFlt3^int^CD9^low^ cells, nor LMPPs generating any megakaryocyte colonies (Figure 7F).

To further improve the efficiency of MegE differentiation, we performed single-cell clonogenic assays in MegE-supporting suspension cultures. In agreement with previous studies (Adolfsson et al. 2005; Mansson et al. 2007), LMPPs lacked detectable MegE potential, whereas a large fraction of clones generated from LSKFlt3^−^ cells contained both megakaryocytes and erythroid cells. Within the LSKFlt3^int^CD9^high^ fraction, 12.46% and 21.3% of the cells produced megakaryocyte and erythroid progeny, respectively. In these conditions, the LSKFlt3^int^CD9^low^ also generated MegE cells, but at a substantially reduced frequency compared to the LSKFlt3^int^CD9^high^ cells (3.3% and 13.8% respectively, Fig. 7G, 7H). In summary, the functional data support identification of a multilineage hematopoietic transition state within the LSKFlt3^int^CD9^high^ cells. This multilineage capacity is rapidly lost as CD9 expression is downregulated (e.g. in Flt3^int^CD9^low^) and Flt3 expression is upregulated (for example in Flt3^high^ LMPP) and paralleled chromatin changes consonant with increasing accessibility of lineage-associated TF motifs.

## DISCUSSION

Advances in scRNA-seq have led to an intense scrutiny, and subsequent revision of the hematopoietic cellular hierarchy by positing the existence of ‘cloud’ HSPCs, where cells exhibit very little distinction in terms of lineage priming while displaying stem-like program (Velten et al. 2017, Giladi A et al. 2018). The cloud in turn, radiates off branches manifesting incremental but progressive gain of lineage-specific gene expression establishing neo-archetypal continuums. However, despite the advocacy of continuums, the origin of such lineage-specific axes within the amorphous cloud state is largely unknown. Moreover, a rigorous pairing of scRNA-seq with lineage tracing has suggested that fate decisions within HSPCs cannot be confidently read out by scRNA-seq alone (Weinreb et al, 2020) despite the possible presence of transitional cell states.

Interestingly, HSPCs arranged along the gradient of Flt3 cell-surface marker expression manifest a somewhat similar contrast between multipotency and lineage segregation, providing an attractive opportunity to dissect lineage restriction. The apparent dichotomy between Flt3^−^ vs. Flt3^+^ LSK compartments is notable: while the Flt3^−^ fraction contains LT-HSCs, ST-HSCs (MPP1), MegE-biased MPP2, and MPP3 populations, the Flt3^+^ pool includes MPP4 progenitors including Flt3^high^ lymphoid-primed LMPPs. Not surprisingly, the multipotency of the Flt3^+^ population within the LSK compartment has been debated (Woolthuis and Park 2016). However, the abrupt separation between Flt3^−^ and Flt3^high^ cells leaves LSKs with an intermediate level of Flt3 (Flt3^int^) as a largely uninvestigated pool of cells that might act as transitional bridge between multipotency and lineage commitment.

Based on the recent observations that the epigenome, rather than the coding transcriptome, more closely aligns with cell potency (Yu VWC. 2017, (Paul et al. 2015) we extensively utilized scATAC-seq to capture the molecular and cellular basis of transition between the cloud of HSPCs, and the continuums of blood and immune cell lineages that emanate from it. Our analyses showed that LT-HSCs could be resolved into two clusters in terms of global chromatin accessibility; cluster 1 being an example of the archetypal HSCs, while cluster 2 bore similarities with cells of the ST-HSCs, and LSKFlt3^int^CD9^high^ cells (Figure 2D). The ST-HSCs, in contrast, appeared to contain an ensemble of cells spread over six clusters suggesting that lineage priming might be seeded at the level of selective chromatin accessibility within chromatin-wise heterogeneous ST-HSCs (Figure 2D). Notably, this is in contrast to the recent scRNA-seq based observations where heterogeneity was not observed within ST-HSC/MPP1 cells (Fraticelli et al 2018).

Although the LSKFlt3^+^ compartment displays a continuum, it is nonetheless heterogeneous, with Flt3^int^ cells emerging as a population positioned between HSCs and Flt3^high^ LMPPs (Figure 1A). Moreover, the LSKFlt3^int^ versus LSKFlt3^high^ cells inhabit distinct parts of the heterogeneity map and exhibit noticeable functional differences, with multipotency restricted to the LSKFlt3^int^CD9^high^ cells (Figure 2D and G, Figure 6-7 and S6). In comparison, the LSKFlt3^int^CD9^low^ cells were similar to the lymphoid-biased Flt3^high^ LMPPs (Figure 2D and G, Figure 6-7 and S6). CD9 is a widely expressed protein from the tetraspanin family, and a cell surface marker for both megakaryocytes and stem cells (Karlsson et al. 2013). Thus, CD9 could be used to capture cells with stem-like molecular program. We also considered the candidature of thrombopoietin receptor MPL, which is shared between MegE and HSCs, and previously shown to enrich for cells with MegE potential in Flt3^+^ cells (Nishikii, Kurita, and Chiba 2017). However, we noted that the majority of Flt3^+^ cells express MPL thereby limiting its efficiency as a marker (Luc et al. 2008). In contrast, high expression of CD9 is found selectively, in about 7% of Flt3^+^ cells making it a better discriminator of cell populations.

The global motif accessibility trend change analysis identified cells and their coincidental chromatin programs at the intersection of primitive and lineage-primed progenitors. Interestingly, this transition state was dominated by the LSKFlt3^int^CD9^high^ cells that emerged apposed to stem cells while also displaying transcriptional imprints of transition to specific lineages (Figure 3-4). Intriguingly, concurrent accessibility of binding motifs of both stem cell-specific as well as Myeloid-Lymphoid and pre-MegE exclusive TFs was commonly observed in this fraction at single-cell level. However, these cells displayed a trend towards lower mean accessibility of stem cell TF motifs than LT and ST-HSCs, but higher mean accessibility of lineage-TF motifs indicating that a subtle but decisive switch in cell-state towards extensive lineage priming is initiated at this stage. In line with this notion, single-cell RNA-seq and multiplexed qPCR signature of the LSKFlt3^int^CD9^high^ cells showed enrichment for genes linked to stem cells, B and T lymphocytes, as well as MegE lineages while CD9^low^ cells in contrast exhibited expression of genes connected to LMPP and pre-GMPs corroborating the findings from scATAC-seq (Figure 5 and 6A-C).

In order to confirm the molecular analysis, we compared LSKFlt3^int^ CD9^high^ and CD9^low^ cells using a combination of in vivo and in vitro assays. Transplantation assays affirmed that CD9^high^ cells are multipotent progenitors with short-term myeloid and long-term lymphoid engraftment potential but with no long-term self-renewal capacity (Figure 6E-F). This fits with the observation that most of the LSKFlt3^int^ CD9^high^ cells are CD150^−^ (Figure S4D) and suggest that these are MPPs (Kiel, Yilmaz, and Morrison 2008). In-vitro culture conditions that promote simultaneous differentiation of several lineages confirmed that LSKFlt3^int^ CD9^high^ cells are more primitive compared to the CD9^low^ by virtue of generating multiple lineages including megakaryocytes and primitive erythroid (BFU-e) progenitor cells (Figure 6-7). However, in cultures designed to promote only erythroid or Meg differentiation, some LSKFlt3^int^ CD9^low^ cells also appeared to have erythroid and Meg potential, albeit with a lower frequency than the CD9^high^ cells. Taken together, the cellular behavior of LSKFlt3^int^ CD9^high^ cells aligns with the findings of a chromatin state and gene expression signature reflecting a concurrent stem cell program and global multilineage priming.

The transition from HSC to frank lineages is multi-faceted, and therefore reliance on a single set of features is unlikely to reveal the full extent of molecular changes. We anticipate that future single cell multi-omics approaches extracting several epigenomic features such as DNA methylation, and chromatin accessibility (Argelaguet et al. 2019) while simultaneously coupled to lineage tracing might yield molecular signatures that more accurately correspond to, temporally precede and hence empower the transition in potency. The scATAC-seq itself provides an elegant solution by utilizing reads mapped to the mitochondrial genome to clonally track the lineages while simultaneously reading out the chromatin features within the same cell (Ludwig et al. 2019; Xu et al. 2019). However, the small HSPC populations analyzed in this study required pooling cells from different mice, and thus clear lineage tracks could not be detected here (data not shown). Alternatively, lineage tracing based on ‘capture and recapture followed by genome sequencing’ as reported recently might be adapted to reconstruct and contrast the clonal relationships emerging from the LSKFlt3^int^ CD9^high^ vs. CD9^low^ cells in future studies (Lee-Six et al. 2018).

Although our work focuses on the Flt3 gradient as a discriminator of multipotency and lineage segregation, recently reported scRNA-seq of LT-HSC and MPP1-4 progenitor subtypes provides an alternative perspective to dissect the same theme (Rodriguez-Fraticelli et al. 2018). Therefore, a comparative analysis of single cell gene expression could reveal to what extent the Flt3^int^CD9^high^ cells are similar to MPP1-4 progenitors, and if a combination of Flt3, CD9, and SLAM markers could further purify transitional cell types. We also note that our analysis of murine HSPCs is broadly consonant with a recent exploration of human hematopoiesis (Buenrostro et al. 2018) in revealing a chromatin basis of continuum yet uncovering substantial heterogeneity within the classic immunophenotypically defined progenitors. However, whether the chromatin continuum of human HSPCs also displays transitional cell states that manifest coincidental stem and multilineage state remains unexplored. Application of change point detection algorithm as done here is likely to inform if such cell states and underlying TF signatures are shared across species. Application of guided gene selection analysis has recently yielded an early lineage segregation signature among human HSC subsets (Belluschi et al. 2018), and identified murine mixed-lineage transitional states (Olsson et al. 2016), however, scATAC-seq provides a more unbiased chromatin wide measurement to discern linage priming likely obviating the need for a supervised analysis.

The intermediary position of LSKFlt3^int^ CD9^high^ cells between the stem and lineage-biased progenitors prompts questions regarding how distinctive cell-states emerge, and the nature of the inductive signals triggering the transition. We foresee that a partial dissolution of stem-like program could stochastically pave way for simultaneous gain of lineage priming within a subset of Flt3^−^ cells that are CD9^+^. Once established, the chromatin program is simply carried forward to and maintained within the CD9^+^ cells of the Flt3^int^ compartment but rapidly vanishes upon loss of CD9 expression. Alternatively, the LSKFlt3^int^ CD9^high^ cells might commence the transitional program de novo without inheriting it from the Flt3^−^ cells; for example, if both CD9 and Flt3 may themselves be involved in receiving, but differentially responding to the induction signals. If so, the ensuing tug of war between the two could precipitate the transition to a cell state where self-renewal is lost but multipotency is still retained.

## ACKNOWLEDGMENTS

We thank Linda Geironson Ulfsson and Ulrich Pfisterer at the Lund University Center for Translational Genomics as well as Anna Fossum, Zhi Ma and Teona Roschupkina at Lund Stem Cell Center FACS core for expert single-cell genomics and flow cytometry, respectively. This work was supported by grants from the Swedish Cancer Society, The Ragnar Söderberg Foundation, the Knut and Alice Wallenberg Foundation, the Swedish Research Council, the Swedish Society for Medical Research, and the Swedish Childhood Cancer Foundation.

## Author contributions

G.K conceived, designed and supervised the study with contributions from C.B and R.K.T; F.S. designed and performed the experiments with contributions from S.W., E.E. C.B., and R.K.T.; P.D. designed and performed the bioinformatics- and computational analyses. F.S, P.D, E.S., C.B., D.B., R.K.T, and G.K. analyzed and interpreted data; F.S., G.K. R.K.T., and P.D. prepared the figures and wrote the manuscript with input from all authors.

## Data and code availability

The data from scATAC-seq, and scRNA-seq have been submitted as GSE14876. All processed data and code required to reproduce the figure from scRNA-seq and scATAC-seq analysis have been deposited at OSF:https://osf.io/y3faj/?view_only=87f2d996e90c49e5a81157e167040c0c

## EXPERIMENTAL PROCEDEURES

### ScATAC-seq and analysis

LSKCD34^−^Flt3^−^ (LT-HSC), LSKCD34^+^Flt3^−^ (ST-HSC), LSKCD34^+^Flt3^int^ (LSKFlt3^int^), LSKCD34^+^Flt3^high^ (LMPP), LSKFlt3^−^CD150^+^CD48^+^ (MPP2), Lin^−^Sca1^−^ Kit^+^FcgRII/III^low^CD150^+^Endoglin^low^CD41^−/low^ (Pre-MegE), LSKCD34^+^Flt3^int^CD9^high^ (LSKCD9^high^), and LSKCD34^+^Flt3^int^CD9^low^ (LSKCD9^low^) cells were sorted, and nuclei isolated following 3 minutes of cell lysis as per low input protocol from 10X Genomics, CA, USA. Single-cell chromatin accessibility profiles were generated by tagmentation followed by library preparation according to the Chromium Single Cell ATAC Reagent Kits User Guide (Satpathy et al. 2019). The libraries were run on TapeStation (Agilent, CA, USA) and sequenced on Nova-Seq (Illumina Inc, CA, USA) using SP100 flow cell with the following specification: 48 (49 for MPP2 and preMegE) cycles for read 1N, 8 cycles for i7 index, 16 cycles for i5 index and 48 (50 for MPP2 and preMegE) cycles for read 2N.

The sequencing data were processed using cellranger-atac pipeline (v 1.1.0), reads aligned to mouse mm10 genome assembly, and only the read-pairs with MAPQ > 20 were retained. MACS2 was used to call peaks with the following specification-s=150 --nomodel --shift=-75 --extsize=150. The data was clustered using the default graph-clustering approach used in Seurat (Satija et al. 2015). The clustering was performed with two values for resolution parameter: 0.75 and 0.1 in order to obtain one fine- and one coarse clustering of the data. Slingshot v 1.2 was used to identify the differentiation trajectories within the data (Street et al. 2018). The FIMO algorithm from MEME suite version 4.12.0 was used to scan the peaks for 571 JASPAR motifs (Fornes et al. 2020). More details on cell identification; peak calling, clustering, motif analysis, overlap with FANTOM enhancers, TF-IDF analysis and motif enrichment within enhancer clusters are provided in supplementary methods.

### Sc-RNAseq and Sc-qPCR data analysis

LSKCD34^+^Flt3^int^CD9^high^, LSKCD34^+^Flt3^int^CD9^low^, and LSKCD34^+^Flt3^int^ cells were sorted from the bone marrow of age-matched C57BL/6J mice, and loaded into 10X Chromium lanes, and the libraries were prepared as per the instruction manual of 10X Chromium. The final libraries were paired-end sequenced using 26 cycles for Read 1, 8 cycles for i7 index, and 98 cycles for read 2.

ScRNAseq data were processed using Cellranger 3.0.2 to generate counts matrices against the mm10 transcriptome. Seurat v2 R package was used to filter and normalize the data. Dimensionality reduction (UMAP) and graph-based clustering was performed using the first fifteen PCA components. Cluster marker genes were identified using Seurat’s default algorithm. The data from BloodSpot ‘normal mouse hematopoiesis’ was used (Bagger, Kinalis, and Rapin 2019) to annotate cell type identity of each cluster. For each cluster, a radar plot was generated using min-max scaled median value of marker genes in each cell type. Nabo v0.3.0 (Parashar Dhapola et al. manuscript in preparation, available here: https://github.com/KarlssonG/nabo) was used to project the cells from LSKFlt3^int^CD9^high^ and LSKFlt3^int^CD9^low^ cells onto the LSKFlt3^int^ graph. Single cell qPCR of index-sorted LSKCD34^+^Flt3^int^ population was performed (Sommarin et al. 2018) and analyzed as described previously (Lang et al. 2015).

### Mice

WT C57Bl/6 (CD45.2) mice were purchased from Taconic. Congenic B6SJL (CD45.1), and C57Bl/6 × B6SJL (CD45.1/CD45.2) mice were kept in ventilated racks and given autoclaved food and water in the barrier facility at the Biomedical Center of Lund University. The ethical committee at Lund University, Sweden approved all mouse experiments.

### Flow cytometry and cell sorting

Peripheral blood, unfractionated BM, or magnetically c-kit-enriched BM from 8- to 11-week-old mice was stained in PBS/fetal calf serum (FCS) for 30 minutes with combinations of antibodies listed in Table S1. Dead cells were excluded using 7-aminoactinomycin D (7-AAD) from Sigma-Aldrich Co, St Louis, MO). Cells were sorted on a FACS-AriaII or AriaIII cell sorter (BD Biosciences, San Jose, CA) or analyzed on a FACSCanto II (BD Biosciences) or BD LSR Fortessa flow cytometer. Analysis performed using FlowJo software (version 10.1r1).

### In vivo analyses (Transplantation)

100 LTHSCs, LSKCD34^+^Flt3^int^CD9^high^ cells, LSKCD34^+^Flt3^int^CD9^low^ cells, and LMPPs, or ten LSKCD34^+^Flt3^int^CD9^high^ cells were FACS sorted and injected intravenously together with 2. 5×10^5^ supportive BM cells (CD45.1/2) into lethally irradiated (900Gy) recipient mice (CD45.1). The CD45.2 donor cells were monitored at 2, 4, 8 and 16 weeks post-transplantation. For secondary transplantations, whole BM was isolated at 16 weeks post-transplant, and 1×10^6^ cells were transplanted into lethally irradiated CD45.1 secondary recipient mice. Donor derived lineage reconstitution was monitored by FACS with antibodies for myeloid (CD11b^+^Gr1^+^) and lymphoid (CD3^+^ B220^+^) cells.

### In vitro analyses

A switch-culture system was set up to evaluate the combined myeloid, B, and erythroid potential from single LTHSCs, LSKFlt3^int^ CD9^high^ cells, LSKFlt3^int^ CD9^low^ cells, and LMPPs. Single cells were seeded onto a 80% confluent OP9 monolayer in 48-well plates containing 100μl medium as described (Nakano, Kodama, and Honjo 1994). Directly after sorting, 100μl medium supplemented with SCF (50 ng/mL), FLT3L (50 ng/mL) was added. After 4 days of co-culture, cells from each of the OP9-containing wells were divided and transferred to either confluent OP9 monolayer wells supplemented with SCF (50 ng/mL), FLT3L (50 ng/mL), IL7 (20 ng/mL), or to SFEM medium with 1% Pen-Strep, 30% FBS, SCF (50 ng/mL), Epo (3 U/mL) and holo-Transferrin (0.3 mg/ml); half of the culture medium was replaced every week. After additional 21 days of co-culture, the cells were FACS analyzed for the presence of Erythroid cells (CD45^−^F4-80^−^Mac1^−^Gr1^−^TER119^+^), B cells (CD45^+^ CD19^+^) and myeloid cells (either CD45^+^ Mac1^+^ Gr1^+^, or CD45^+^ Mac1^+^ F-480^+^). In addition, the clones were required to have >50-gated events of surface markers within scatter profile to be counted as positive.

For expansion in liquid culture, 500 cells were sorted per well in 96-well plates and were grown in SFEM StemSpan supplemented with 30% FBS (StemCell Technologies Inc. Canada) as described in (Pietras et al. 2015). Evaluation of Meg and Erythroid lineage potential at single cell level was performed as previously described with minor modifications (Adolfsson et al. 2005).

### Statistical analysis

In vivo and vitro assays were analyzed using GraphPad Prism 7 as indicated in the figure legends. The statistical analyses for single cell ATAC, mRNA-seq is provided in the respective methods.

## SUPPLEMENTARY METHODS

Supplementary methods include detailed description of scATAC-seq, scRNA-seq, sc-qPCR analysis and in vitro assays and are provided separately.

