## Supplementary methods, figures and tables for "Concurrent stem- and lineage-affiliated chromatin programs precede hematopoietic lineage restriction"

#### ScATAC-seq analysis

##### *Cell identification*

The sequencing data were processed using cellranger-atac pipeline (v 1.1.0) from 10x Genomics to obtain the aligned reads. The pipeline annotates each aligned read with a cell barcodes (corrected or uncorrected). Mouse mm10 genomic assembly was used as the reference. The number of fragments associated with each barcode was used as a filter to remove the background barcodes. Barcode multiplets were removed using 10x Genomics recommended scripts for v1.1.0. Additionally, the cells with percentage mitochondrial contamination > 3 or less than -1 (both values in log2 scale) were removed.

##### *Peak calling*

Fragments associated with valid cell barcodes was obtained from cellranger-atac generated BAM files. Only properly aligned read-pairs, each with MAPQ > 20 were retained. Based on the read pairs, both ends of the fragments (accounting for soft clipping of reads during alignment) were identified and these were considered as cut sites and used in downstream analysis. A cell-barcode was allowed only one cut site for a particular genomic location. The cut sites were aggregated from all the cell barcodes, de-duplicated and saved in BED format where each line is 1 nucleotide-long range with start position indicating the cut site. MACS2 was called on the aggregated BED files separately for each chromosome to speed up the peak calling process. Default MACS2 parameters were overridden with -s=150 --nomodel --shift=-75 --extsize=150 and the genome size parameter was set as the respective chromosome size. The narrow peaks from each chromosome were merged to create a reference peak set for use in the downstream analyses. For each cell, the number of cut sites within the peak set was counted, and the resulting cell-peak matrix was saved in the MTX format.

*Clustering, low dimensional embedding and trajectory identification*

The MTX files containing peak-cell matrices for each sample were loaded in Seurat v 3.0.1 and concatenated. Only cells that had total cut sites within the range of 25,000-500,000 were retained. The data was log-normalized and the 20,000 most variable peaks were identified using 'vst' method. PCA was run on the variable features. UMAP was run to obtain a low dimensional embedding of the data for the top six principal components, and 20 neighbours were used as parameters for UMAP. The data was clustered using the default graph-clustering approach used in Seurat (Satija et al. 2015). The clustering was performed with two values for resolution parameter: 0.75 and 0.1 in order to obtain fine- and coarse clustering of the data. The differentiation trajectories within the data were identified using Slingshot v 1.2 (Street et al. 2018). Slingshot was provided fine clustering identity of each cell, and the trajectory was set to start from the cluster at a terminal end of the UMAP containing LT-HSCs. The cells with weight <1 for a trajectory were excluded for that particular trajectory.

*Motif identification*

The total peak-set was used to identify occurrences of transcription factor binding site motifs within peaks. We used the FIMO algorithm from MEME suite version 4.12.0. Each JASPAR motif ((Fornes et al. 2020) was scanned through the peaks individually. For each motif, the peaks were refined with p-value <0.05, giving a peak-set for each motif. The peak-set was divided in proximal or distal set based on peak's genomic location. Peaks within -2000 to +500 region of transcription start sites (TSS) were considered as 'proximal' peaks. The peaks that were neither proximal, nor within gene bodies were considered as 'distal' peaks. Thereafter, two cell-motif matrices were created; one for the proximal, and the other for the 'distal' peak sets wherein each column is a motif and each row is a cell. Each column in the cell-motif matrix is the rowsum of subset of the columns (peaks for a given motif) of cell-matrix.

*Cell-Motif analysis*

The cell-motif matrix is normalized by dividing each column's values by column sum, and multiplying by a scaling factor of 1000, and calculating z-scores for each motif. To calculate cluster-cluster correlation, proximal or distal matrix was processed to calculate median values of cells within each cluster; and the resultant cluster-motif matrix was used to calculate Pearson's r (as implemented in scipy package) value between each clusters. Cell-cell correlation was calculated by using Pearson's r-values, and the resultant matrix was clustered using 'ward' linkage method as implemented in the scipy package. The dendrogram of the clustered cell-cell matrix was cut into the same number of branches as the number of fine clusters as detected by Seurat in complete peak-set data. The concordance between Seurat clusters and motif clusters was calculated using Homogeneity metric as implemented in the scikit-learn package. To identify enriched motifs in each of the coarse clusters, Mann-Whitney U test as implemented in mannwhitneyu function of stats module of scipy package was used, and -Log10 (P values) were min-max transformed to obtain the relative p-values. Only those motifs with variance to mean ratio below (or equal to) 0.025 were used. This was done to make sure that motif prevalence is not restricted to small subset of a cluster. To produce pseudotime profiles of transcription factor binding motifs, the motif accessibility of the cells was smoothened using centered rolling mean over the window of 200 cells. Ward agglomerative clustering was performed on the TFBS-pseudotime profiles, and the dendrogram was cut to get 20 clusters for both Myeloid-Lymphoid and MegE trajectories. The TFBS clusters were ordered in the heatmap based on the position of highest mean accessibility along the pseudotime. To identify the trend change positions along the pseudotime, the window sliding method from the ruptures package was used to identify one change point in each TFBS pseudotime-ordered profile. The change point positions were aggregated across all TFBS profile to find the trend-change regions in the pseudotime.

*Overlap with FANTOM mouse enhancers*

The promoter-distal peaks were investigated to identify enhancers. Permissive enhancer coordinates for mouse genome were obtained from FANTOM5 project and were converted from mm9 to mm10 genomic coordinates using UCSC's liftover tool. Since, the enhancers were identified using CAGE method, only the expressed portion of the enhancer is captured. These enhancer coordinates were extended by 1Kb on both side and intersected with the distal peaks. This identified 12,438 putative enhancers regions in the dataset.

*TF-IDF normalization methodology*

To perform TF-IDF normalization, a matrix of accessibility values (raw number of cut fragments within peaks) was created for all the cells and enhancer peaks. Total enhancer fragments were calculated (by summation) for each cell and the values were divided by the respective total value. This provided the TF values (term frequency) of each enhancer in each cell. To calculate IDF (inverse document frequency), the cell number divided by the sum of TF values for each enhancer and log2 transformation was applied. Finally, products of TF and IDF values were calculated to obtain normalized values. The matrix of normalized values was subsetted for either My-Ly cells or MegE cells. The z-scores for individual enhancer were calculated before visualization as heat map. The 20 partitions of enhancers were created using hierarchical clustering allowing identification of groups that show altered accessibility along the pseudotime.

*Motif enrichment within enhancers*

For a given subset of enhancer peaks, we identified the number of peaks that contain motif for each TFBS. The same is also done for the rest of enhancers that are not present in the subset (negative subset from 12,438 enhancers). To identify the enrichment of the motif presence in the subset, Fisher's exact test is performed. The p-values for all the TFBS motifs are corrected using Benjamini-Hochberg at 5% false discovery rate and reported. To summarize the enhancer accessibility for cell clusters following steps were performed: a subset of enhancers was selected for example, a particular cluster of enhancers from pseudotime trajectory. The total accessibility values (TF-IDF normalized) for each enhancer was transformed to z-score and summation of all enhancers values was calculated for each cell so that there is a singular value for each cell. These values were visualized in boxplots by partitioning cells into cell clusters.

**ScRNAseq analysis**

Cell-cycle effect was removed during the scaling step by regressing against G2M and S scores that were generated by ‘Cell Cycle Score’ function. Dimensionality reduction was done using UMAP and graph-based clustering was performed using the first fifteen PCA components. Cluster marker genes were identified using Seurat’s default algorithm. To annotate cell type identity of each cluster, data from BloodSpot ‘normal mouse hematopoiesis’ was used (Bagger, Kinalis, and Rapin 2019). For each cluster, a radar plot was generated using min-max scaled median value of marker genes in each cell type.

Nabo v0.3.0 (*Parashar Dhapola et al. manuscript in preparation; available here:* https://github.com/KarlssonG/nabo) was used for cell projection analysis; the cells from LSKFlt3$^{int}$CD9$^{high}$ and LSKFlt3$^{int}$CD9$^{low}$ cells were projected onto the LSKFlt3$^{int}$ graph using modified Canberra metric. All the projected cells were classified into Seurat defined clusters using Nabo’s ‘Graph.classify_target’ function.

**Sc-qPCR and data Analysis**

Multiplexed quantitative real-time PCR analyses (BioMark 48.48 or 96.96 Dynamic Array platform (Fluidigm)) with Taqman Gene Expression Assays listed in Table S2 (Applied Biosystems, CA, USA) were performed on index-sorted cells from LSKCD34$^{+}$Flt3$^{int}$ population as described previously (Sommarin et al. 2018). The data was analyzed using the single Cell Expression Visualizer web tool (Lang et al. 2015).

**In vitro analyses**

For colony assay, 150 cells from LSKFlt3$^{int}$ CD9$^{high}$ or CD91$^{ow}$ populations sorted and were plated in 35-mm petri dish in Methylcellulose (GM M3434). Cells were incubated at 37°C in 5% CO2. Total colony number was scored after 12-14 days of culture.

For CFU-Mk assay, 250 LTHSCs, LSKCD34$^{+}$Flt3$^{int}$CD9$^{high}$ cells, and LSKCD34$^{+}$Flt3$^{int}$CD9$^{low}$ cells were plated in MegaCult (Catolog#04900) according to the manufacturer’s instruction (Stem Cell Technologies Inc.).

**SUPPLEMENTARY FIGURES**

**Figure S1**. (Related to Fig.1)

Representative FACS gating strategy and sort purity check for **(A)** LTHSCs, STHSCs, LSKFlt3$^{int}$, LMPP, LSKFlt3$^{int}$CD9$^{high}$, LSKFlt3$^{int}$CD9$^{low}$ and **(B)** MPP2 and Pre-MegE populations.

**Figure S2**. (Related to Figure 1 and 2)

Left panel: ScATAC-seq fragment size distribution for each sorted HSPC population, Right panel: Relative enrichment around TSS for each population

**Figure S3.** (Related to Figure 1 and 2)

ScATAC-seq cell barcode identification from each of the 8 HSPC populations, Left panel: Inverted frequency plot in Log10 showing number of fragments associated with each barcode. The plot shows how the number of fragments associated with each barcode decreases as we consider increasing number of low frequency barcodes. The first drop in the curve is expected to mark the true cell barcodes from background barcodes. The number along the vertical line marks the number of barcodes that were considered as cell barcodes. In the colored area, the percentage indicates the number of fragments that were present in cell barcodes. Middle panel: Frequency versus (fragments/cell) for cell barcodes and a few top non -cell barcodes, Right panel: fragment numbers from cells plotted against the % mitochondrial read content. The cells shown in blue were removed from the analysis on account of either too high or too low mitochondrial content. The cells shown in black were further removed by cellranger ad hoc script provided by 10x genomics to remove doublets/multiplets.

**Figure S4.** (Related to Fig.2 and Fig.3)

(A-C) Examples of UMAP plots depicting the TFBSs accessibility of selected TFs in either distal or promoter proximal regions of each cell. The dark red colour indicates high accessibility of TFBS and dark blue indicates low accessibility. **(D)** Representative FACS plot showing the CD9$^{high}$ cells within the LSKFlt3$^{int}$ fraction and expression of SLAM markers in LT-HSCs, ST-HSCs, LSKCD34$^{+}$Flt3$^{int}$ and LMPP, **(E)** UMAP plot highlighting individual populations. **(F)** Cluster heat map showing correlation values (Pearson's r values) between each pair of fine clusters. Correlations were based on merge accessibility of motif carouse promoter proximal region.

**Figure S5.** (Related to Fig.5)

**(A)** The heat map shows the molecular signature of selected genes across clusters 1 to 10 as observed in scRNA-seq. **(B)** Lineage annotation of clusters. Cells from each cluster have been highlighted in the UMAP and also in the DDRTree layout of the cells. The contour plot over DDRTree depicts the density of cells from each cluster. The radar plot shows the normalized median expression of top 20 cluster marker genes in different hematopoietic cell types.

**Figure S6. In vivo reconstitution from limiting doses of LSKFlt3$^{int}$CD9$^{high}$ cells (**Related to Fig.6)

**(A)** Donor reconstitution in peripheral blood of recipient mice transplanted with 10 LSKFlt3$^{int}$CD9$^{high}$ cells. Each dot represents one recipient mouse (n=3). **(B)** Representative FACS-plots of donor contribution to the LSK compartment in BM of recipient mice following sixteen-week transplantation of 10 LSKFlt3$^{int}$CD9$^{high}$ cells compared to the c-Kit FMO control. **(C)** Mean donor lineage distribution over time in recipients transplanted with 10 LSKFlt3$^{int}$CD9$^{high}$ cells. The error bars represent standard deviation (n=3 biological experiment).

**Figure S7. The LSKFlt3$^{int}$ CD9 $^{high}$ cells are multipotent (**Related to Fig.6)

Representative FACS-plots showing the fraction of LSK cells **(A)** (n=3) and myeloid cells **(B)** (n=2) 3, 6 and 9 days following culture in myeloid differentiation-culture of 500 cells from each indicated LSK population.

#### SUPPLEMENTARY TABLES

Table S1: Antibodies used for surface immunophenotyping and sorting for hematopoietic stem and progenitor cells

Table S2: Single cell multiplexed qPCR: genes and corresponding commercially available Taqman probes (related to figure 5)

Table S3: Transcription factors belonging to different clusters along the Lympho-Myeloid pseudotime

Table S4: Transcription factors belonging to different clusters along the MegE pseudotime

S-Figure 1

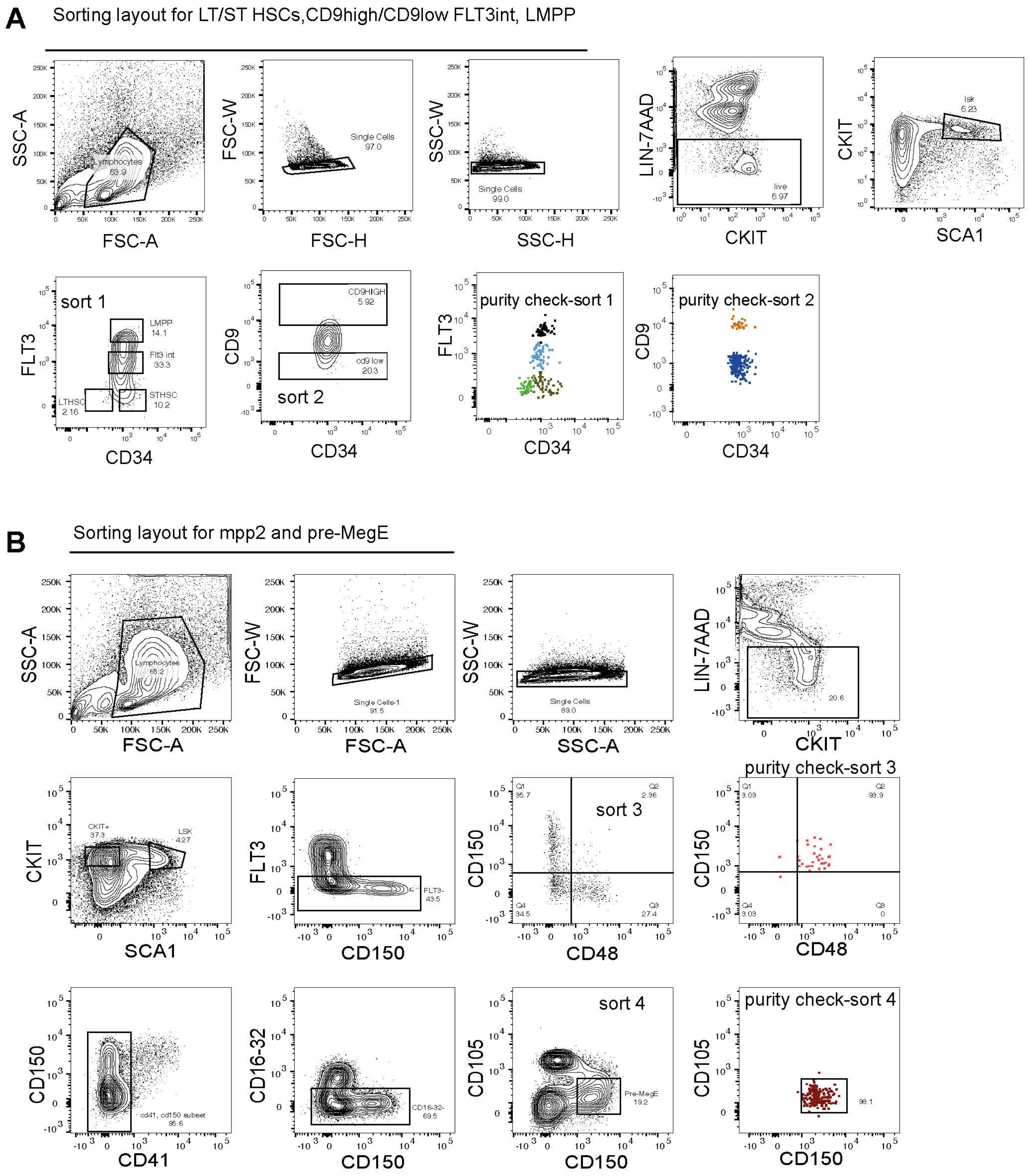
A
Sorting layout for LT/ST HSCs,CD9high/CD9low FLT3int, LMPP
SSC-A
FSC-A
Lymphocytes 63.9
FSC-W
FSC-H
Single Cells 97.0
SSC-W
SSC-H
Single Cells 99.0
LIN-7AAD
CKIT
live 6.97
CKIT
SCA1
lsk 6.23
sort 1
FLT3
CD34
LMPP 14.1
Flt3 int 33.3
STHSC 10.2
LTHSC 2.16
CD9
CD34
CD9HIGH 5.92
cd9 low 20.3
sort 2
purity check-sort 1
FLT3
CD34
purity check-sort 2
CD9
CD34
B
Sorting layout for mpp2 and pre-MegE
SSC-A
FSC-A
Lymphocytes 65.2
FSC-W
FSC-A
Single Cells-1 91.5
SSC-W
SSC-A
Single Cells 89.0
LIN-7AAD
CKIT
20.6
CKIT
SCA1
CKIT+ 37.3
LSK 4.27
FLT3
CD150
FLT3- 43.5
CD150
CD48
sort 3
Q1 35.7
Q2 2.36
Q3 27.4
Q4 34.5
purity check-sort 3
CD150
CD48
Q1 3.03
Q2 93.9
Q3 0
Q4 3.03
CD150
CD41
cd41, cd150 subset 95.6
CD16-32
CD150
CD16-32- 69.5
CD105
CD150
sort 4
Pre-MegE 19.2
purity check-sort 4
CD105
CD150
98.1

S-Figure 2

A

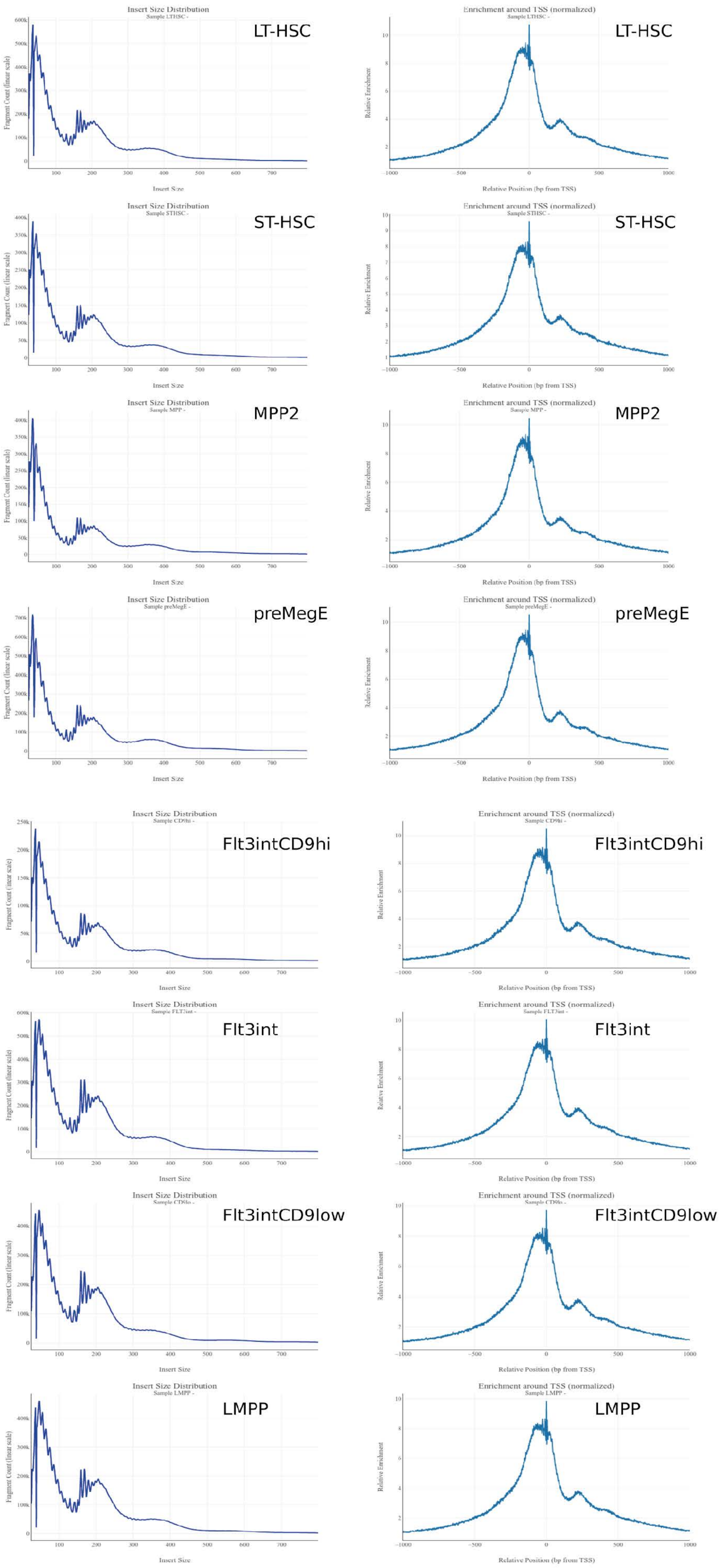

Insert Size Distribution
Fragment Count (linear scale)
Insert Size
Enrichment around TSS (normalized)
Relative Enrichment
Relative Position (bp from TSS)
LT-HSC
ST-HSC
MPP2
preMegE
Flt3intCD9hi
Flt3int
Flt3intCD9low
LMPP

### S-Figure 3

A

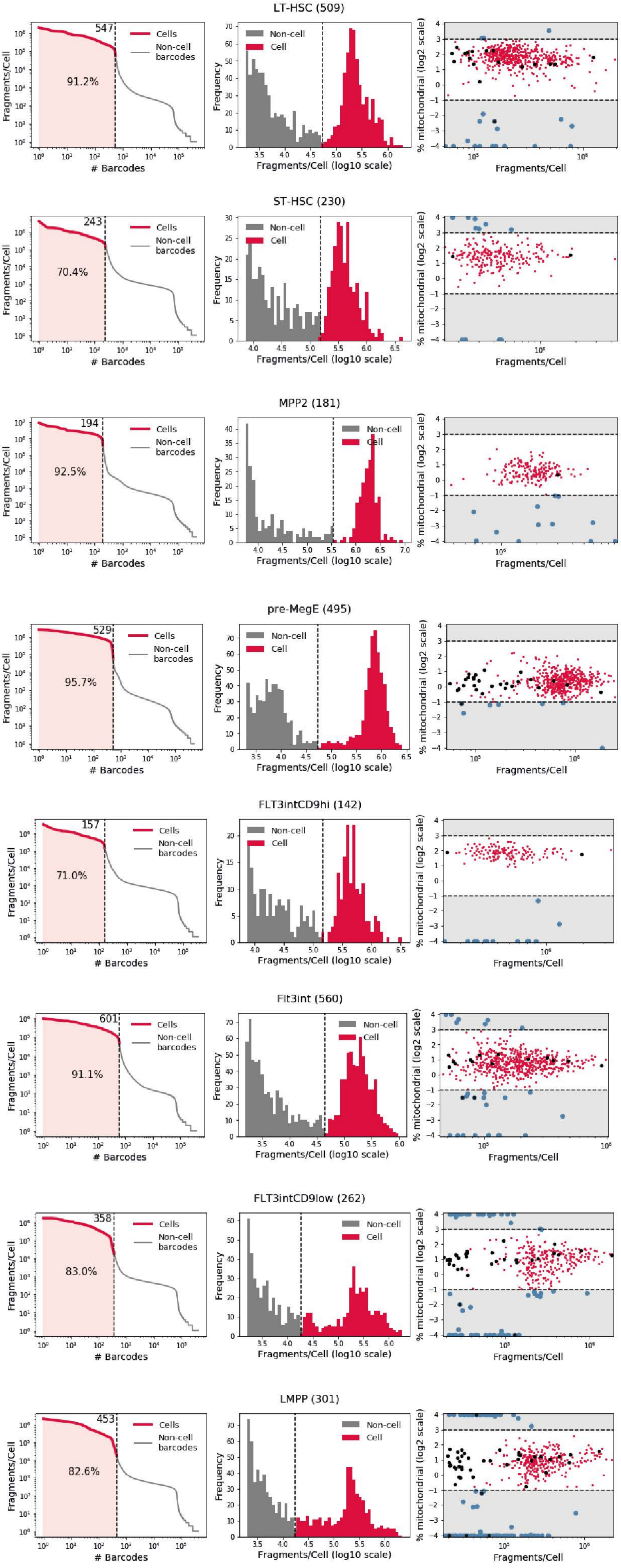
LT-HSC (509)
547
91.2%
Cells
Non-cell barcodes
Non-cell
Cell
Fragments/Cell
### Barcodes
Frequency
Fragments/Cell (log10 scale)
% mitochondrial (log2 scale)
ST-HSC (230)
243
70.4%
MPP2 (181)
194
92.5%
pre-MegE (495)
529
95.7%
FLT3intCD9hi (142)
157
71.0%
Flt3int (560)
601
91.1%
FLT3intCD9low (262)
358
83.0%
LMPP (301)
453
82.6%

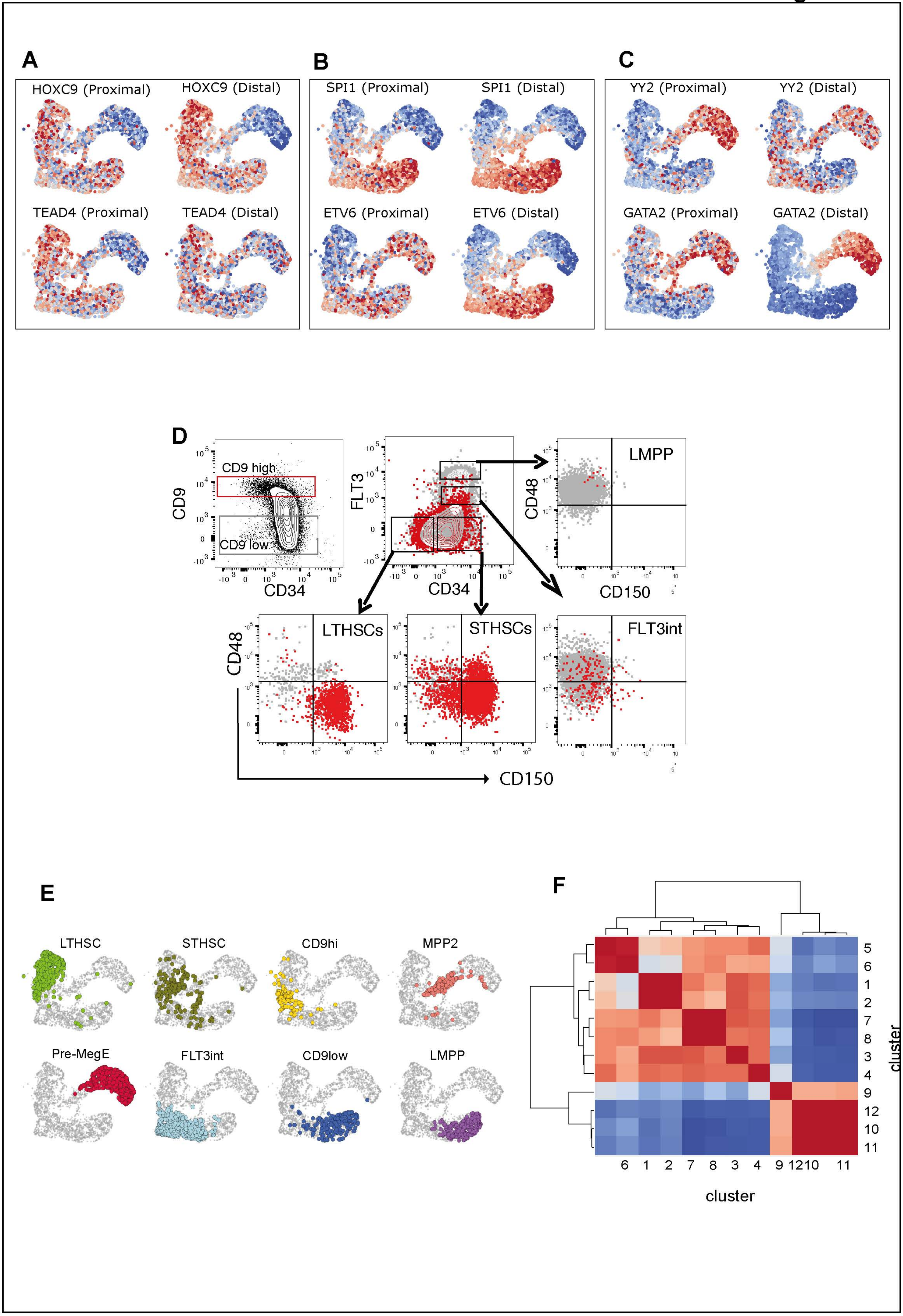
A
HOXC9 (Proximal)
HOXC9 (Distal)
TEAD4 (Proximal)
TEAD4 (Distal)
B
SPI1 (Proximal)
SPI1 (Distal)
ETV6 (Proximal)
ETV6 (Distal)
C
YY2 (Proximal)
YY2 (Distal)
GATA2 (Proximal)
GATA2 (Distal)
D
CD9 high
CD9 low
CD9
CD34
FLT3
CD34
LMPP
CD48
CD150
LTHSCs
STHSCs
FLT3int
CD48
CD150
E
LTHSC
STHSC
CD9hi
MPP2
Pre-MegE
FLT3int
CD9low
LMPP
F
5
6
1
2
7
8
3
4
9
12
10
11
cluster
6 1 2 7 8 3 4 9 12 10 11
cluster

S-Figure 5

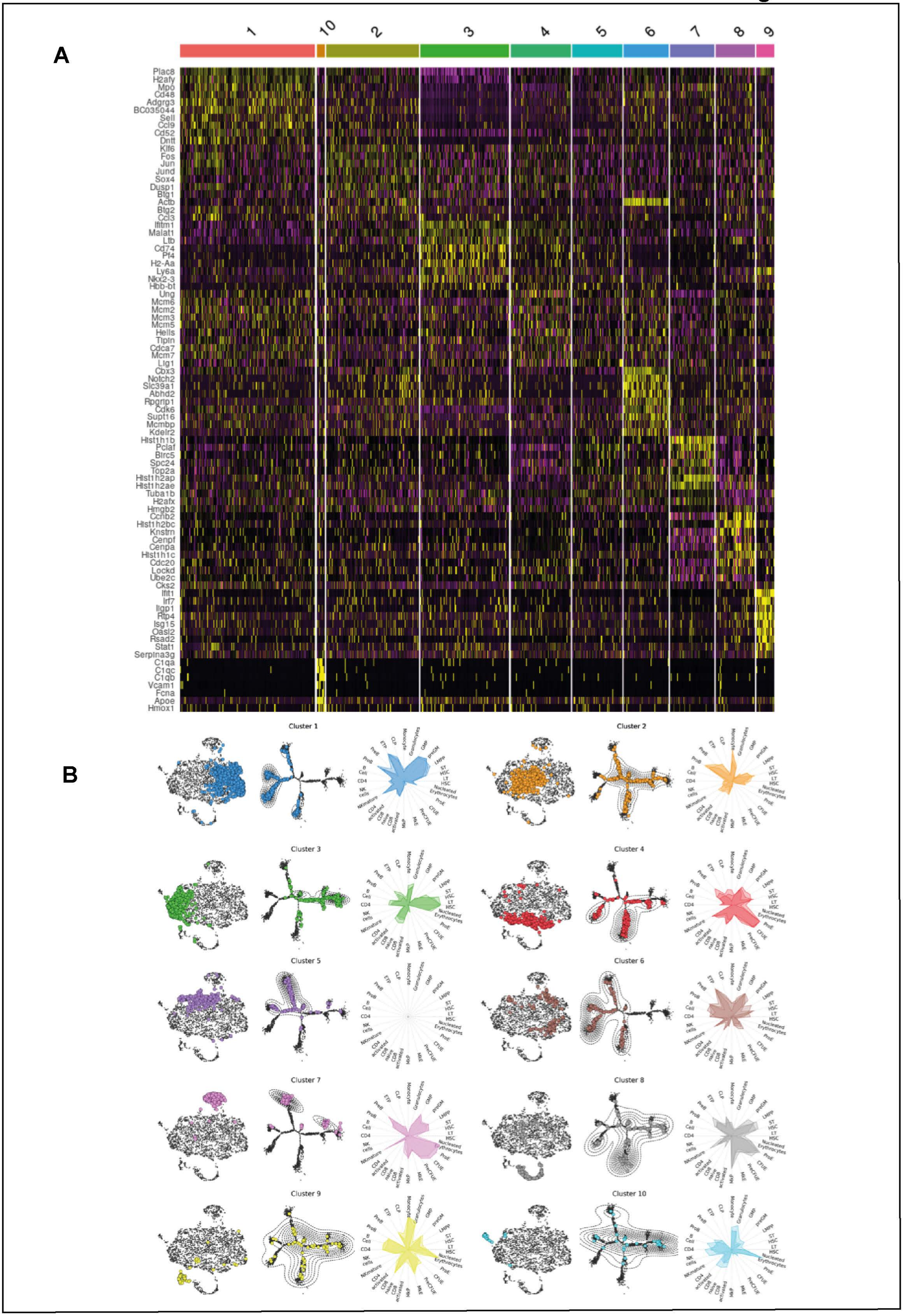
A
1
10
2
3
4
5
6
7
8
9
Plac8
H2afy
Mpo
Cd48
Adgrg3
BC035044
Sell
Ccl9
Cd52
Dntt
Klf6
Fos
Jun
Jund
Sox4
Dusp1
Btg1
Actb
Btg2
Ccl3
Ifitm1
Malat1
Ltb
Cd74
Pf4
H2-Aa
Ly6a
Nkx2-3
Hbb-bt
Ung
Mcm6
Mcm2
Mcm3
Mcm5
Hells
Tipin
Cdca7
Mcm7
Lig1
Cbx3
Notch2
Slc39a1
Abhd2
Rpgrip1
Cdk6
Supt16
Mcmbp
Kdelr2
Hist1h1b
Pclaf
Birc5
Spc24
Top2a
Hist1h2ap
Hist1h2ae
Tuba1b
H2afx
Hmgb2
Ccnb2
Hist1h2bc
Knstrn
Cenpf
Cenpa
Hist1h1c
Cdc20
Lockd
Ube2c
Cks2
Ifit1
Irf7
Iigp1
Rtp4
Isg15
Oasl2
Rsad2
Stat1
Serpina3g
C1qa
C1qc
C1qb
Vcam1
Fcna
Apoe
Hmox1
B
Cluster 1
Cluster 2
Cluster 3
Cluster 4
Cluster 5
Cluster 6
Cluster 7
Cluster 8
Cluster 9
Cluster 10

S-Figure 6

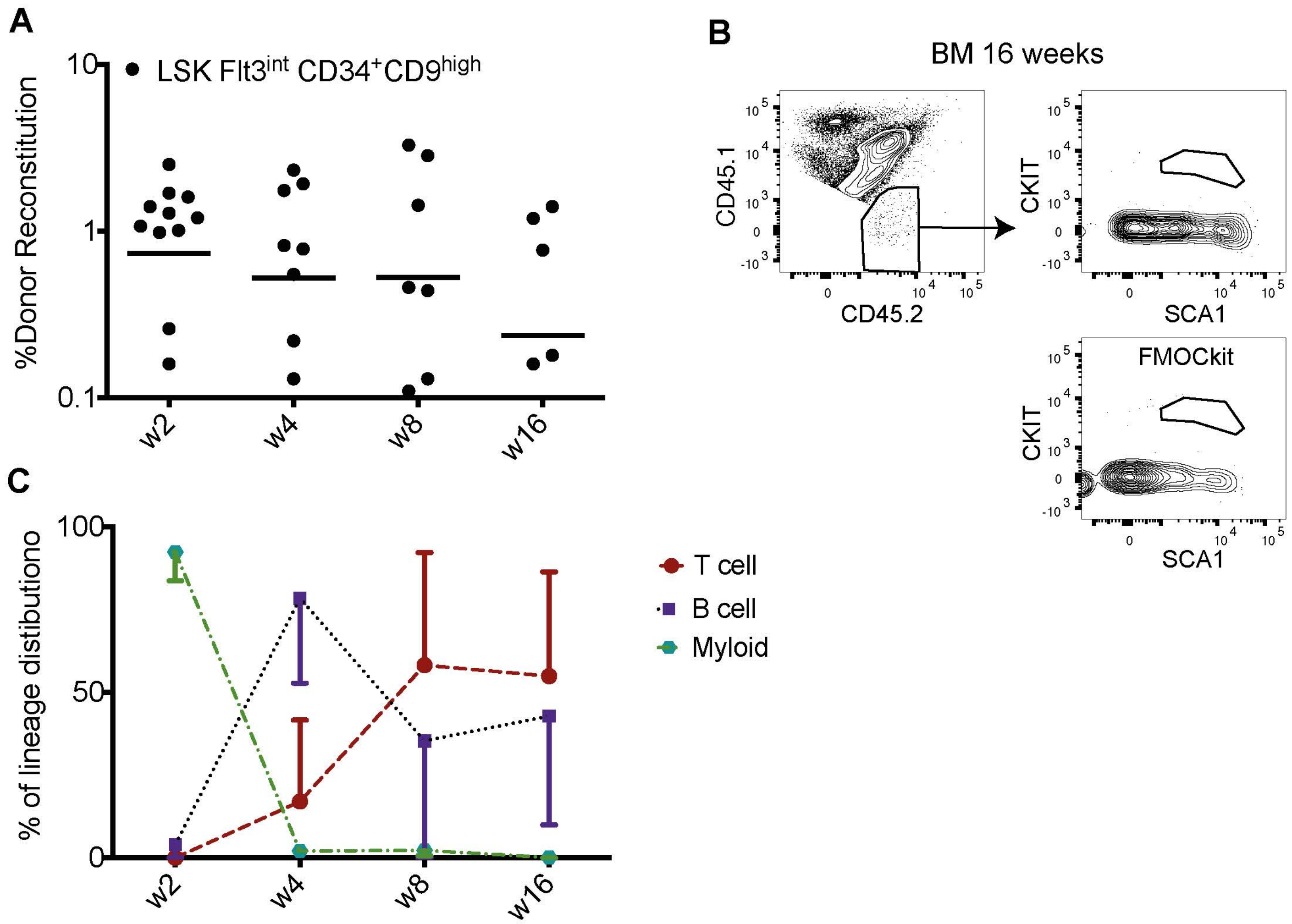

A
LSK Flt3int CD34+CD9high
%Donor Reconstitution
10
1
0.1
w2
w4
w8
w16
B
BM 16 weeks
CD45.1
CD45.2
CKIT
SCA1
FMOCkit
CKIT
SCA1
C
% of lineage distibutiono
100
50
0
w2
w4
w8
w16
T cell
B cell
Myloid

S-Figure 7

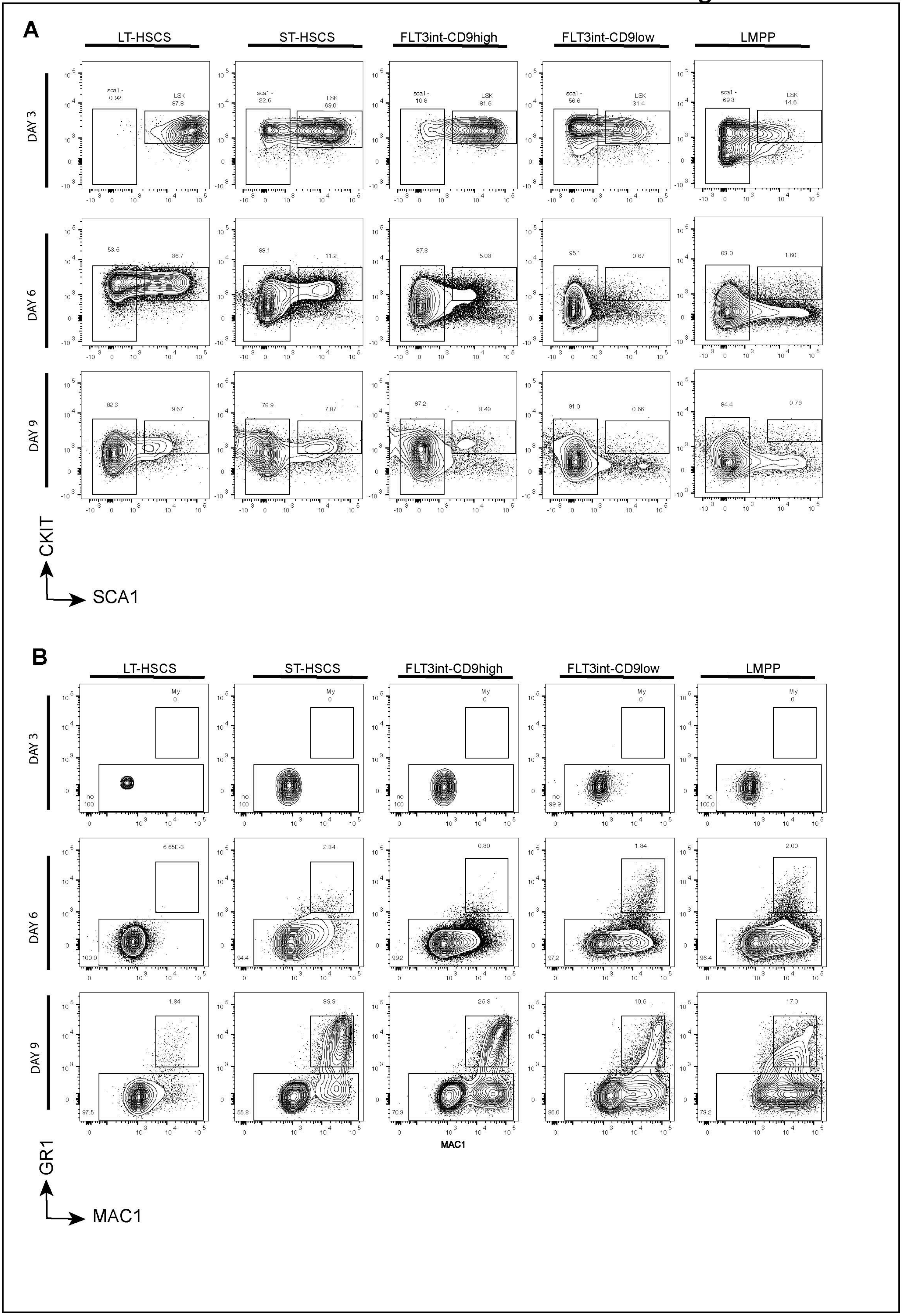

A
LT-HSCS
ST-HSCS
FLT3int-CD9high
FLT3int-CD9low
LMPP
DAY 3
DAY 6
DAY 9
sca1 -
0.92
LSK
87.8
22.6
69.0
10.8
81.6
56.6
31.4
69.3
14.6
53.5
36.7
83.1
11.2
87.3
5.03
95.1
0.87
83.8
1.60
82.3
9.67
78.9
7.87
87.2
3.48
91.0
0.66
84.4
0.78
CKIT
SCA1
B
LT-HSCS
ST-HSCS
FLT3int-CD9high
FLT3int-CD9low
LMPP
DAY 3
DAY 6
DAY 9
My
0
no
100
99.9
100.0
6.65E-3
2.34
0.30
1.84
2.00
94.4
99.2
97.2
96.4
1.84
39.9
25.8
10.6
17.0
97.5
55.8
70.3
86.0
73.2
MAC1
GR1
MAC1

SUPPLEMENTAL TABLES

| Antibody | Fluorochrome | Clone | Company |
|---|---|---|---|
| Sca-1 | BV-421 | D7 | Biolegend |
| Sca-1 | PE-CY7 | D7 | Biolegend |
| CKIT | APC | 2B8 | Biolegend |
| CD135 (Flt3) | PE | A2F10.1 | BD Biosciences |
| CD135 (Flt3) | BV-421 | A2F10.1 | BD Biosciences |
| CD34 | FITC | RAM34 | eBioscience |
| CD34 | Biotin | RAM34 | eBioscience |
| CD9 | PE | MZ3 | Biolegend |
| CD9 | FITC | MZ3 | Biolegend |
| CD9 | Biotin | MZ3 | Biolegend |
| CD150 | BV605 | TC15-12F12.2 | Biolegend |
| CD150 | PE-CY7 | TC15-12F12.2 | Biolegend |
| CD48 | APC-CY7 | HM48-1 | Biolegend |
| CD41 | BV-711 | MWReg30 | BD Biosciences |
| CD41 | PE | MWReg30 | BD Biosciences |
| CD16/CD32 | Alexa Fluor 700 | 93 | eBioscience |
| CD105 | PE-CY7 | MJ7/18 | Biolegend |
| CD11b | APC | M1/70 | Biolegend |
| CD11b | PE-CY5 | M1/70 | Biolegend |
| CD11b | PE | M1/70 | Biolegend |
| GR-1 | BV-605 | RB6-8C5 | Biolegend |
| GR-1 | PE-CY5 | RB6-8C5 | Biolegend |
| GR-1 | APC-CY7 | RB6-8C5 | Biolegend |
| GR-1 | PE | RB6-8C5 | Biolegend |
| F4/80 | APC-CY7 | BM8 | Biolegend |
| CD45R/B220 | PE-CY5 | RA3-6B2 | Biolegend |
| CD45R/B220 | PE | RA3-6B2 | Biolegend |
| CD45R/B220 | APC-CY7 | RA3-6B2 | Biolegend |
| CD3 | PE-CY5 | 145-2C11 | Biolegend |
| TER-119 | PE-CY5 | TER-119 | Biolegend |
| TER-119 | PE-CY7 | TER-119 | Biolegend |
| NK-1.1 | PE-CY5 | PK136 | Biolegend |
| NK-1.1 | BV-421 | PK136 | Biolegend |
| NK-1.1 | PE | PK136 | Biolegend |
| CD45 | Alexa Fluor 700 | 30-F11 | Biolegend |
| CD45.2 | APC | 104 | Biolegend |
| CD45.1 | PE-CY7 | A20 | Biolegend |
| CD45.1 | FITC | A20 | Biolegend |
| CD19 | PE-CY7 | eBio1D3 | eBioscience |
| MPL | Biotin | No10403 | IBL |

**Tabel S1**-FACS antibodies, related to Experimental procedure.

| Gene symbol | Assay ID | Gene symbol | Assay ID |
|---|---|---|---|
| Gata2 | Mm00492301_m1 | Il7r | Mm00434295_m1 |
| MPL | Mm00440310_m1 | Rag1 | Mm01270936_m1 |
| Procr | Mm00440992_m1 | Rag2 | Mm01270938_m1 |
| Pf4 | Mm00451315_g1 | Hes1 | Mm01342805_m1 |
| Pbx1 | Mm04207617_m1 | Ccr9 | Mm02620030_s1 |
| Cited2 | Mm01188099_g1 | Ccr7 | Mm01301785_m1 |
| Vwf | Mm00550376_m1 | Gata3 | Mm00484683_m1 |
| Apoe | Mm01307193_g1 | Il2ra | Mm01340213_m1 |
| Meis1 | Mm00487664_m1 | Foxo1 | Mm00490671_m1 |
| Cd48 | Mm00455932_m1 | Satb1 | Mm01268940_m1 |
| *Hlf* | Mm00723157_m1 | Mpeg1 | Mm01222137_g1 |
| Slamf1/CD150 | Mm00443316_m1 | Runx1 | Mm01213404_m1 |
| Laptm4b | Mm00835799_g1 | Lfng | Mm00456128_m1 |
| Hey1 | Mm00468865_m1 | Ikzf1 | Mm01187877_m1 |
| Kit | Mm00445212_m1 | PU.1/Spi1 | Mm00488140_m1 |
| Fgd5 | Mm00554954_m1 | Id2 | Mm00711781_m1 |
| Tie1 | Mm00441786_m1 | Pou6f1 | Mm00447791_m1 |
| CD120-B/Tnfrsf1b | Mm00441889_m1 | Thy1 | Mm00493681_m1 |
| Cd184/Cxcr4 | Mm01996749_s1 | Dtx1 | Mm00492297_m1 |
| Cd34 | Mm00519283_m1 | CSF2RA | Mm00438331_g1 |
| Notch1 | Mm00627185_m1 | LEF1 | Mm00550265_m1 |
| Flt3 | Mm00439016_m1 | Ebf1 | Mm00432954_m1 |
| Xpc | Mm01183434_m1 | Ick | Mm00517387_m1 |
| Ctsg | Mm00456011_m1 | Runx3 | Mm00490666_m1 |
| Hhex | Mm00433954_m1 | Dntt | Mm00493500_m1 |
| Cd200 | Mm00487740_m1 | Tcf7 | Mm00493445_m1 |
| Ly6d | Mm00521959_m1 | Irf8 | Mm00492567_m1 |
| Cd244 | Mm01296079_m1 | Lck | Mm00802897_m1 |
| Cd8a | Mm01182107_g1 | Cd123/Il3ra | Mm00434273_m1 |
| Mpo | Mm01298424_m1 | Epor | Mm00833882_m1 |
| Csf1r/Cd115 | Mm01266652_m1 | Klf1 | Mm00516096_m1 |
| Csf2rb | Mm00655745_m1 | Car2 | Mm00501576_m1 |
| Fcgr3 | Mm00438882_m1 | Phf10 | Mm00470370_m1 |
| Ly86 | Mm00440240_m1 | Cd41/Itga2b | Mm00439741_m1 |
| Cebpa | Mm00514283_s1 | Nfe2 | Mm00801891_m1 |
| Fes | Mm01318102_m1 | GP1ba | Mm00501677_g1 |
| Spen | Mm00465639_m1 | Cited4 | Mm00550568_s1 |
| Anxa1 | Mm00440225_m1 | Gata1 | Mm01352636_m1 |
| Lyz2 | Mm01612741_m1 | Gfi1b | Mm00492318_m1 |
| Lyz1 | Mm00657323_m1 | Cd9 | Mm00514275_g1 |
| Csf3r/Cd114 | Mm00432735_m1 | Tal1 | Mm01187033_m1 |
| Klf4 | Mm00516104_m1 | Ccna2 | Mm00438063_m1 |
| Gfi1 | Mm00515853_m1 | Ccnb2 | Mm01171453_m1 |
| Fli1 | Mm00484410_m1 | Ccnd1 | Mm00432359_m1 |
| Ccne1 | Mm01266311_m1 | Mki67 | Mm01278617_m1 |

| | | | |
|---|---|---|---|
| Ccnf | Mm00432385_m1 | Atp5a1 | Mm00431960_m1 |
| p21/Cdkn1 | Mm00432448_m1 | Hprt | Mm01545399_m1 |
| Cdkn1b | Mm00438168_m1 | | |

**Table S2**- Taqman assay-Related experimental procedure in Figure 5,
Genes expressed at low-variation across the data set as well as genes with undetectable expression was excluded from Figure 5

**Table 3**

| 1 | 2 | 3 | 4 | 5 | 6 | 7 | 8 | 9 | 10 | 11 | 12 | 13 | 14 | 15 | 16 | 17 | 18 | 19 | 20 |
|---|---|---|---|---|---|---|---|---|---|---|---|---|---|---|---|---|---|---|---|
| JUND | LHX3 | HOXB3 | MYBL1 | GATA1 | SIX2 | TCFL5 | IRF9 | HOXA11 | HOXC13 | RARG | SP1 | ZNF423 | THAP1 | USF1 | PRDM1 | FOSL2::JUN_VA | TEAD1 | POU2F3 | BHLHA15 |
| FOS | CUX2 | EVX1 | POU4F1 | GATA3 | POU3F2 | HEY1 | IRF8 | TFCP2 | JUNB_VAR.2 | PAX6 | GABPA | BCL6B | RFX3 | TP73 | RORA_VAR.2 | FOSB::JUNB_VA | ID4 | ONECUT2 | ZBTB18 |
| FOSL2::JUND | DUX | GRHL1 | HOXA5 | NR3C2 | POU5F1B | ALX3 | IRF1 | HOXD11 | SIX3 | HMX2 | ZIC4 | NR2C2 | PAX1 | KLF1 | MEF2C | CREB5 | TCF3 | POU5F1 | MEIS2 |
| NFE2 | HOXD12 | HOXB13 | MEOX2 | NOTO | NR3C1 | ZBTB33 | IRF3 | HOXC10 | TEF | SOX8 | E2F1 | GSX1 | ESR2 | NR4A2 | SMAD3 | RARA | TBX2 | POU2F2 | JUN::JUNB_VAR.2 |
| FOS::JUNB | HOXC9 | NKX2-5 | NFATC3 | IRF5 | POU3F1 | HINFP | IRF4 | | PROX1 | MITF | CLOCK | TFEB | ELF4 | SOX2 | MEF2D | JDP2_VAR.2 | NFKB1 | CEBPB | ETV2 |
| FOSL1::JUNB | FOXJ3 | EMX2 | NFATC1 | PRRX2 | POU3F4 | LHX8 | RARB | | DMBX1 | NRL | E2F6 | KLF12 | HIC2 | BATF3 | CDX1 | FOSL2::JUNB_V | NR4A1 | POU1F1 | MYOG |
| STAT1::STAT2 | MAF::NFE2 | FOXC1 | SOX11 | LHX6 | POU2F1 | ISX | | | DBP | SREBF2_VAR | MLXIPL | RARG_VAR.2 | GCM1 | TCF7L1 | CDX2 | CREB1 | RORA | SOX15 | OLIG3 |
| JUN::JUNB | PAX3 | IRF7 | NFATC2 | VENTX | | FOXD3 | | | FOSL1::JUN_ | NR1H3::RXR | CTCF | RREB1 | RXRG | CREB3 | | FOS::JUN_VAR. | CREB3L1 | CEBPD | TP63 |
| FOSL1::JUND | FOXJ2 | HOXB5 | NFAT5 | BSX | | MTF1 | | | FOSB::JUN | MAFB | ELK3 | HNF4G | NR2F6_VAR.2 | ZNF143 | | | MGA | DUXA | TFAP4 |
| MEF2B | FOXL1 | ARID3B | PHOX2A | GBX2 | | RBPJ | | | FOSL2::JUND | NFIL3 | ATF3 | GLIS1 | KLF13 | NHLH1 | | | TBX5 | CEBPE | BHLHE22 |
| FOSL2::JUNB | FOXO3 | HMX1 | TEAD4 | ATF1 | | RAX2 | | | | SREBF1_VAR | ARNT::HIF1A | CENPB | NR4A2::RXRA | NR2F1 | | | SCRT2 | CEBPG | REL |
| JUNB | LIN54 | STAT1 | HOXB2 | EMX1 | | FOXC2 | | | | ZBED1 | HES7 | TFAP2C_VAR.2 | FEV | ELF3 | | | TFEC | ARID5A | HSF4 |
| BACH2 | HOXD13 | MAFG | TEAD2 | HNF1A | | E2F3 | | | | | TFAP2C | XBP1 | RFX5 | NKX3-1 | | | TBX21 | | DUX4 |
| JDP2 | FOXI1 | MNX1 | DLX1 | EN2 | | BHLHE40 | | | | | SP8 | HNF4A | NR1A4::RXRA | EHF | | | POU4F2 | | ASCL2 |
| POU3F3 | FOXD2 | VAX2 | TEAD3 | LBX1 | | HOXC12 | | | | | ETV4 | PPARA::RXRA | RFX1 | KLF4 | | | EOMES | | ASCL1 |
| MEF2A | FOXA1 | HMX3 | RORC | GATA5 | | BHLHE41 | | | | | CTCFL | INSM1 | RFX2 | YY1 | | | TBR1 | | PKNOX2 |
| FOSL2::JUN | GRHL2 | DLX2 | SOX9 | HLTF | | LHX9 | | | | | TFAP2B_VAR.2 | SIX1 | ZBTB7C | ERG | | | NR2F2 | | TGIF1 |
| FOS::JUN | MAFF | NOBOX | SOX4 | BCL6 | | EGR1 | | | | | OTX2 | ZBTB7B | RXRB | NKX2-8 | | | NEUROD1 | | TWIST1 |
| FOSL1::JUN | FOXP1 | BARHL2 | PHOX2B | LMX1B | | MAX | | | | | TFAP2A_VAR.3 | GLIS3 | GLI2 | GFI1 | | | NFKB2 | | NKX3-2 |
| FOSB::JUNB | FOXO1 | SOX21 | | E2F2 | | SP2 | | | | | RHOXF1 | CREB3L2 | ELK4 | RARA::RXRG | | | TBX1 | | CEBPA |
| FOSL1 | MAFK | NKX6-1 | | LMX1A | | PAX2 | | | | | ZNF740 | PAX9 | VDR | FLI1 | | | TBX20 | | SOX1 |
| FOSL2 | FOXP3 | ISL2 | | EN1 | | MXI1 | | | | | ETV3 | YY2 | ESRRG | TCF7L2 | | | SNAI2 | | PKNOX1 |
| FOS::JUND | BACH1::MAFK | POU6F2 | | HOXD3 | | ATF7 | | | | | KLF14 | E2F8 | ZNF282 | DDIT3::CEBPA | | | SPIC | | RUNX1 |
| ONECUT1 | HOXA10 | GSC | | UNCX | | EGR2 | | | | | PAX5 | PPARG::RXRA | ZNF263 | AR | | | MYB | | SMAD2::SMAD3::SMAD4 |
| JUN_VAR.2 | FOXQ1 | GATA1::TAL1 | | HESX1 | | HES1 | | | | | TFAP2A_VAR.2 | EBF1 | SOX3 | REST | | | NKX2-5_VAR.2 | | NEUROD2 |
| BATF::JUN | NFIA | FOXB1 | | LHX4 | | EGR4 | | | | | SPDEF | NR2F6 | ELF1 | KLF9 | | | ZNF410 | | TAL1::TCF3 |
| | CUX1 | VSX2 | | RAX | | NRF1 | | | | | TFAP2C_VAR.3 | RFX4 | SPZ1 | EWSR1-FLI1 | | | SCRT1 | | MYF6 |
| | GSC2 | POU4F3 | | BARX1 | | CREM | | | | | ELK1 | NR5A2 | | NKX2-3 | | | SOX10 | | DMRT3 |
| | FOXK2 | PROP1 | | PRRX1 | | NFIC::TLX1 | | | | | TFAP2A | MZF1_VAR.2 | | ZEB1 | | | TBP | | HSF2 |
| | PAX7 | VAX1 | | ALX4 | | EGR3 | | | | | ZIC3 | RXRA | | RARA_VAR.2 | | | ETV6 | | MECOM |
| | DLX4 | IRF2 | | GATA2 | | GMEB2 | | | | | TFAP2B_VAR.3 | ESRRA | | SRF | | | TBX4 | | PBX2 |
| | FOXO6 | VSX1 | | LHX2 | | NFYB | | | | | GMEB1 | NR1H4 | | ETS1 | | | SREBF2 | | MSC |
| | ONECUT3 | FOSL1::JUND_VAR.2 | | NKX6-2 | | GCM2 | | | | | E2F7 | MAX::MYC | | HIC1 | | | TWIST2 | | OLIG2 |
| | SRY | | | ESX1 | | MSX3 | | | | | SP4 | | | SOX6 | | | GFI1B | | JUN |
| | FOXO4 | | | MSX1 | | HAND1::TCF3 | | | | | TFE3 | | | LEF1 | | | HLF | | NEUROG2 |
| | HOXD9 | | | STAT4 | | ZNF24 | | | | | ZIC1 | | | ESRRB | | | ESR1 | | ATF4 |
| | FOXD1 | | | STAT3 | | NPAS2 | | | | | NFYA | | | MYBL2 | | | TCF7 | | RELA |
| | HOXA13 | | | ZNF384 | | MYC | | | | | ETV5 | | | | | | TBX15 | | MEIS3 |
| | NFE2L2 | | | RELB | | HIF1A | | | | | OTX1 | | | | | | HMBOX1 | | TP53 |
| | FOXF2 | | | EVX2 | | BARHL1 | | | | | KLF5 | | | | | | PPARG | | RUNX2 |
| | DLX3 | | | MIX-A | | E2F4 | | | | | SP3 | | | | | | TCF4 | | SOX17 |
| | DLX6 | | | POU6F1 | | ZFX | | | | | USF2 | | | | | | SPI1 | | TCF21 |
| | HOXA9 | | | MIXL1 | | GLIS2 | | | | | TFAP2B | | | | | | ELF5 | | ATOH1 |
| | FOXG1 | | | STAT5A::STAT5B | | HEY2 | | | | | ARNTL | | | | | | FIGLA | | SMAD4 |
| | FOXP2 | | | ALX1 | | NFIX | | | | | PLAG1 | | | | | | FOXH1 | | RHOX11 |
| | FOXK1 | | | GATA4 | | MNT | | | | | NR1H2::RXRA | | | | | | | | OLIG1 |
| | FOXA2 | | | STAT6 | | MLXIP | | | | | PITX1 | | | | | | | | MYOD1 |
| | | | | PDX1 | | HOXC11 | | | | | MLX | | | | | | | | PBX1 |
| | | | | MEOX1 | | LBX2 | | | | | ETV1 | | | | | | | | POU5F1::SOX2 |
| | | | | GBX1 | | TFDP1 | | | | | KLF16 | | | | | | | | RXRA::VDR |
| | | | | GATA6 | | MYCN | | | | | ID2 | | | | | | | | NR2E1 |
| | | | | HNF1B | | HOXA2 | | | | | ZBTB7A | | | | | | | | BHLHE23 |
| | | | | GSX2 | | SHOX | | | | | PITX3 | | | | | | | | T |
| | | | | CRX | | ARX | | | | | | | | | | | | | SOX13 |
| | | | | MSX2 | | HES5 | | | | | | | | | | | | | TBX19 |
| | | | | | | SHOX2 | | | | | | | | | | | | | ZSCAN4 |
| | | | | | | HOXD8 | | | | | | | | | | | | | SREBF1 |
| | | | | | | PAX4 | | | | | | | | | | | | | MEIS1 |
| | | | | | | NFIC | | | | | | | | | | | | | NR2E3 |
| | | | | | | HES2 | | | | | | | | | | | | | RARB_VAR.2 |
| | | | | | | | | | | | | | | | | | | | RUNX3 |
| | | | | | | | | | | | | | | | | | | | TCF12 |
| | | | | | | | | | | | | | | | | | | | JUND_VAR.2 |
| | | | | | | | | | | | | | | | | | | | NEUROG1 |
| | | | | | | | | | | | | | | | | | | | ERF |
| | | | | | | | | | | | | | | | | | | | TGIF2 |
| | | | | | | | | | | | | | | | | | | | PBX3 |
| | | | | | | | | | | | | | | | | | | | RORB |
| | | | | | | | | | | | | | | | | | | | RARA::RXRA |
| | | | | | | | | | | | | | | | | | | | HSF1 |

**Tabel 4**

| 1 | 2 | 3 | 4 | 5 | 6 | 7 | 8 | 9 | 10 | 11 | 12 | 13 | 14 | 15 | 16 | 17 | 18 | 19 | 20 |
|---|---|---|---|---|---|---|---|---|---|---|---|---|---|---|---|---|---|---|---|
| CEBPD | MAFG | HMX1 | HOXB2 | SOX3 | HSF1 | GLIS1 | TCF4 | TBX5 | NR2E3 | RFX3 | EMX1 | PBX1 | LHX3 | USF1 | HOXD8 | HIF1A | MYBL2 | DLX1 | HES5 |
| RAX | FOXJ3 | FOSL1::JUNB | ESR1 | MTF1 | HNF1B | TFAP2C | TBR1 | SCRT1 | SREBF2 | OLIG3 | LHX2 | MEF2A | POU6F1 | FOSL2::JUNB_VAR.2 | RUNX2 | SOX15 | YY2 | TP63 | PAX2 |
| NFATC1 | NFE2L2 | FOSL1::JUND_VAR.2 | PAX9 | GLIS2 | TCF7L2 | NR4A2::RXRA | RORB | ID4 | SPIC | TAL1::TCF3 | LBX1 | MEF2C | LIN54 | ETV5 | CRX | HOXC13 | KLF9 | LHX8 | NRF1 |
| MAFK | VSX1 | GSC | NFIA | BARHL1 | FOSL2::JUN_VAR.2 | ZBTB7B | NR4A1 | REL | PPARG::RXRA | FOXH1 | CDX2 | GFI1B | HOXA5 | PBX3 | BHLHE22 | SOX8 | RARG_VAR.2 | TP73 | ZBTB33 |
| DLX3 | HOXD11 | FOS | NFIC | MLXIPL | TEF | GCM1 | NRL | FIGLA | RORA | RXRB | DUXA | HSF2 | MEOX2 | CREB5 | ZNF740 | HEY2 | POU2F1 | NFYA | TFCP2 |
| FOXP2 | PDX1 | JUNB | RAX2 | RARB_VAR.2 | ERF | PAX5 | HOXA2 | GBX1 | NR2F6 | ELF4 | ARID5A | FOSB::JUN | SOX11 | ETV3 | OLIG1 | ID2 | ZIC4 | SCRT2 | HES2 |
| CEBPB | SRY | ALX4 | PPARG | GRHL2 | STAT3 | GLIS3 | TBX1 | ZEB1 | HOXD9 | SPDEF | VENTX | ELK1 | MIX-A | PBX2 | LHX6 | CREM | KLF1 | NFKB1 | BHLHE40 |
| HOXC11 | FOXQ1 | FOSL1::JUN | CUX1 | AR | TEAD3 | TFAP2B | TBX21 | MLX | EHF | HIC2 | | GLI2 | HOXC10 | GSX1 | PROX1 | ZBED1 | GMEB1 | PAX4 | POU5F1B |
| HOXC12 | MAFF | SOX21 | UNCX | ESR2 | IRF3 | TFAP2A_VAR.2 | TBX15 | TBX4 | SREBF1 | PKNOX2 | | FOSB::JUNB_VAR.2 | RELB | RFX1 | TWIST2 | ESX1 | TFE3 | NFKB2 | EGR1 |
| FOXB1 | MAF::NFE2 | NFE2 | MSX3 | RARB | SOX4 | | SIX1 | RELA | E2F8 | RFX4 | | JUN::JUNB_VAR.2 | PAX7 | TGIF1 | POU1F1 | NR3C2 | POU2F3 | ATF1 | BHLHE41 |
| GBX2 | VSX2 | FOS::JUND | TFAP2B_VAR.2 | | HAND1::TCF3 | | MYOG | SPZ1 | RARA_VAR.2 | RFX5 | | NR1H2::RXRA | | TGIF2 | RREB1 | BARHL2 | PITX1 | MYF6 | MAX |
| NOBOX | BACH2 | FOSL1::JUND | DLX6 | | GSC2 | | NR2F2 | MSC | RARA::RXRA | OLIG2 | | MEF2D | | TBP | SRF | SOX9 | PHOX2A | TP53 | HES1 |
| IRF7 | | BARX1 | BSX | | ELF5 | | EN2 | SNAI2 | TBX20 | SOX17 | | MEF2B | | PKNOX1 | CEBPA | MLXIP | ZIC3 | | ATF3 |
| HOXB3 | | FOS::JUN | MSX1 | | SMAD3 | | TCF21 | TCF3 | THAP1 | HLF | | FOS::JUN_VAR.2 | | HMBOX1 | NKX2-5_VAR.2 | FOSL1::JUN_VAR.2 | CREB3L1 | | POU4F2 |
| NKX6-1 | | RORC | EMX2 | | EVX1 | | ASCL2 | | EOMES | ZBTB18 | | ISL2 | | RUNX1 | DMRT3 | POU5F1::SOX2 | SP3 | | POU3F4 |
| NFATC3 | | FOSL1 | NKX6-2 | | GABPA | | HNF4G | | NKX2-3 | PPARA::RXRA | | ATF7 | | ETV4 | ATOH1 | CLOCK | POU3F3 | | ONECUT1 |
| GRHL1 | | NR3C1 | NR1A4::RXRA | | RXRA::VDR | | ESRRA | | FEV | RXRA | | ELK3 | | RFX2 | MEIS1 | ARNTL | POU2F2 | | FOXD3 |
| FOXK1 | | ARX | MSX2 | | PRDM1 | | MYOD1 | | TCF12 | TWIST1 | | | | BCL6B | SOX1 | | GATA2 | | HEY1 |
| FOXO3 | | ZFX | CUX2 | | NKX2-5 | | CDX1 | | SPI1 | ELF1 | | | | PLAG1 | YY1 | | MECOM | | MXI1 |
| FOXP3 | | BCL6 | ISX | | NR2E1 | | RBPJ | | RARA::RXRG | RHOXF1 | | | | | GFI1 | | GATA6 | | TFEB |
| FOXD1 | | STAT5A::STAT5B | HOXA11 | | ETS1 | | T | | SOX2 | | | | | | NEUROD1 | | CREB3 | | LHX4 |
| VAX2 | | IRF4 | STAT1::STAT2 | | POU6F2 | | ESRRG | | ZNF282 | | | | | | HMX3 | | E2F3 | | ALX3 |
| SOX10 | | GCM2 | LBX2 | | FLI1 | | ASCL1 | | SOX13 | | | | | | NEUROG2 | | CREB3L2 | | CTCFL |
| FOXL1 | | JDP2 | NKX3-2 | | ALX1 | | NOTO | | RXRG | | | | | | ZSCAN4 | | OTX2 | | ARNT::HIF1A |
| DLX2 | | IRF8 | NKX3-1 | | HSF4 | | NFIC::TLX1 | | ETV6 | | | | | | HMX2 | | NFYB | | LMX1B |
| FOXG1 | | TEAD4 | SHOX | | NR2C2 | | ESRRB | | TCF7 | | | | | | ZNF143 | | MYB | | POU4F3 |
| HESX1 | | FOSL2::JUNB | NHLH1 | | DBP | | MGA | | ATF4 | | | | | | REST | | SP1 | | TCFL5 |
| DLX4 | | FOSL2::JUND | PRRX1 | | ELK4 | | TBX2 | | LEF1 | | | | | | RUNX3 | | RHOX11 | | NR1H3::RXRA |
| FOXO4 | | IRF2 | TBX19 | | RORA_VAR.2 | | NR1H4 | | | | | | | | EVX2 | | ZIC1 | | HINFP |
| FOXK2 | | FOSB::JUNB | SHOX2 | | FOSL2::JUND_VAR.2 | | SMAD4 | | | | | | | | TFEC | | GATA1::TAL1 | | E2F7 |
| IRF5 | | BATF::JUN | TFAP2A | | TCF7L1 | | | | | | | | | | E2F6 | | GATA4 | | JUNB_VAR.2 |
| FOXI1 | | FOXA2 | SREBF2_VAR.2 | | INSM1 | | | | | | | | | | MEIS3 | | RARA | | ONECUT3 |
| FOXP1 | | JUN::JUNB | EBF1 | | HNF1A | | | | | | | | | | MZF1_VAR.2 | | POU3F1 | | E2F1 |
| EN1 | | FOSL2::JUN | HOXA10 | | ETV2 | | | | | | | | | | KLF13 | | ZNF384 | | NPAS2 |
| CEBPG | | STAT1 | PAX1 | | E2F2 | | | | | | | | | | VDR | | POU3F2 | | HES7 |
| FOXO6 | | IRF9 | MAFB | | HOXD3 | | | | | | | | | | ETV1 | | PHOX2B | | CTCF |
| FOXO1 | | FOSL2 | NFIX | | NR2F1 | | | | | | | | | | NEUROG1 | | KLF16 | | MYCN |
| CEBPE | | TEAD2 | PRRX2 | | HOXA9 | | | | | | | | | | ARID3B | | CENPB | | E2F4 |
| FOXF2 | | NFAT5 | TFAP2C_VAR.2 | | TEAD1 | | | | | | | | | | TFAP4 | | SP8 | | POU4F1 |
| HOXD12 | | JUND | GMEB2 | | NR4A2 | | | | | | | | | | PAX6 | | SP4 | | MYBL1 |
| HOXB13 | | IRF1 | LHX9 | | NFIL3 | | | | | | | | | | ZNF263 | | ZBTB7C | | ZNF24 |
| HOXD13 | | NFATC2 | SREBF1_VAR.2 | | STAT4 | | | | | | | | | | BHLHE23 | | GATA5 | | TFDP1 |
| SIX3 | | GSX2 | | | SIX2 | | | | | | | | | | MEIS2 | | TFAP2B_VAR.3 | | LMX1A |
| HOXA13 | | FOXA1 | | | MITF | | | | | | | | | | BATF3 | | USF2 | | MYC |
| VAX1 | | FOS::JUNB | | | ZBTB7A | | | | | | | | | | MEOX1 | | TFAP2A_VAR.3 | | MNT |
| MNX1 | | JUN_VAR.2 | | | JUN | | | | | | | | | | DUX4 | | RARG | | ONECUT2 |
| FOXJ2 | | | | | SMAD2::SMAD3::SMAD4 | | | | | | | | | | ZNF410 | | HLTF | | EGR3 |
| BACH1::MAFK | | | | | DUX | | | | | | | | | | NR5A2 | | FOXC1 | | EGR4 |
| MIXL1 | | | | | CREB1 | | | | | | | | | | SOX6 | | KLF4 | | POU5F1 |
| HOXB5 | | | | | ELF3 | | | | | | | | | | NEUROD2 | | PITX3 | | FOXC2 |
| FOXD2 | | | | | HOXC9 | | | | | | | | | | ZNF423 | | KLF12 | | EGR2 |
| | | | | | STAT6 | | | | | | | | | | EWSR1-FLI1 | | KLF14 | | XBP1 |
| | | | | | JDP2_VAR.2 | | | | | | | | | | HNF4A | | TFAP2C_VAR.3 | | |
| | | | | | NKX2-8 | | | | | | | | | | DDIT3::CEBPA | | KLF5 | | |
| | | | | | JUND_VAR.2 | | | | | | | | | | DMBX1 | | OTX1 | | |
| | | | | | ERG | | | | | | | | | | BHLHA15 | | NR2F6_VAR.2 | | |
| | | | | | HIC1 | | | | | | | | | | | | MAX::MYC | | |
| | | | | | PAX3 | | | | | | | | | | | | PROP1 | | |
| | | | | | | | | | | | | | | | | | SP2 | | |
| | | | | | | | | | | | | | | | | | GATA3 | | |
| | | | | | | | | | | | | | | | | | GATA1 | | |
